# Prediction of Stimulation-Defined Eloquent Cortex Using Graph-Theoretical Connectivity from Electrocorticography During Presurgical Mapping

**DOI:** 10.64898/2026.05.01.722255

**Authors:** Nikasadat Emami, Andrew Michalak, Amirhossein Khalilian-Gourtani, Antoine Ratouchniak, Chenqian Le, Xupeng Chen, Orrin Devinsky, Werner K Doyle, Patricia Dugan, Daniel Friedman, Yao Wang, Adeen Flinker

## Abstract

**Background and Objectives:** Electrical stimulation mapping (ESM) is the clinical gold standard for identifying eloquent cortex during presurgical evaluation but is time-intensive, constrained by incomplete cortical sampling, and limited by patient tolerance. We investigated whether features derived from electrocorticography (ECoG), including functional connectivity measures, can provide complementary information for identifying functionally critical cortex and examined how predictive performance varies across functional domains, behavioral tasks, and data quantity.

**Methods:** Fourteen patients undergoing intracranial monitoring for epilepsy surgery performed speech production tasks while ECoG was recorded independently of stimulation mapping. Graph-theoretic functional connectivity features derived from high-gamma activity (70–150 Hz), combined with anatomical region encoding, were used to train machine learning classifiers, and predictive performance was evaluated using leave-one-subject-out validation with ESM-defined functional deficits used as ground-truth outcomes.

**Results:** Fourteen patients were included. Performance differed across functional domains, with motor-critical electrodes identified with the highest accuracy (ROC–AUC 0.929 ± 0.061; PR–AUC 0.755 ± 0.191), followed by speech arrest (ROC–AUC 0.793 ± 0.103; PR–AUC 0.550 ± 0.196), whereas language-critical electrodes were more difficult to robustly predict (ROC–AUC 0.761 ± 0.160; PR–AUC 0.385 ± 0.167). Additional signal-derived features provided limited benefit beyond anatomical and connectivity features. Performance also varied across tasks and with the number of available trials, with motor prediction remaining stable across trial counts and speech arrest prediction improving with increasing trial counts up to approximately 10 trials before plateauing. In addition, evaluation at the level of stimulation pairs increased sensitivity, with only modest changes in overall performance.

**Discussion:** Graph-theoretic connectivity analysis of ECoG provides complementary information for identifying stimulation-defined functional criticality and supports presurgical mapping. Differences in predictive performance across functional domains likely reflect underlying neuro-physiologic organization, with connectivity providing the clearest improvement for speech arrest prediction. Combined-label prediction strategies increased sensitivity at the expense of specificity, reflecting a clinically relevant tradeoff between broad detection of critical cortex and precise localization. These findings suggest that connectivity-informed approaches may help guide task selection and improve mapping efficiency while complementing electrical stimulation mapping.

## 1 Introduction

Patients with drug-resistant epilepsy are indicated for resective brain surgery to remove the seizure onset zone [1, 2]. However, resection of functionally critical cortex can result in permanent impairments in language or motor function. Presurgical functional mapping is therefore essential to avoid causing functional deficits.

Electrical stimulation mapping (ESM) remains the clinical standard for identifying eloquent cortex during presurgical evaluation [3, 4]. By transiently disrupting cortical function via brief trains of electrical current, ESM can induce behavioral deficits and identify eloquent cortex, reducing the risk of postoperative neurological impairment. However, ESM is time-intensive, depends on patient participation and tolerance, and is constrained by incomplete electrode sampling and practical limitations of testing multiple electrode pairs and functional domains [3]. Approaches using passively recorded neural activity to identify functionally critical cortex may overcome these limitations [5–10].

Subdural electrocorticography (ECoG) provides direct recordings of cortical activity with high temporal and spatial resolution and has been widely used to study speech and language function [7, 10, 11]. Task-related high-gamma activity (70–150 Hz) increases reliably during speech production and yields a robust electrophysiological marker of cortical engagement [12, 13], providing an opportunity to examine neural activity associated with speech and language function. Prior work has applied machine learning to these signals to predict stimulation-defined outcomes [14], but these approaches focus on local electrode activity and do not account for network interactions. Network-based analyses demonstrate that functional interactions among distributed cortical regions are critical for cognition [15–18], and language-critical sites identified by ESM can span multiple subnetworks [19]. These observations suggest that task-related connectivity patterns may provide complementary information to ESM for identifying functionally critical cortex.

In this study, we developed and evaluated a graph-theoretic, connectivity-based framework to estimate the likelihood that individual cortical sites are functionally critical using ECoG signals recorded during speech tasks performed independently of stimulation mapping. Neural activity was analyzed at the trial level and aggregated to generate electrode-level probability estimates. The framework integrates anatomical location with graph-theoretic measures derived from high-gamma functional connectivity and was evaluated using leave-one-subject-out validation against stimulation mapping outcomes.

Using this approach, we addressed three clinically relevant questions. First, we examined how predictive performance varies across speech arrest, motor control, language function, and combined functions. Second, we evaluated which behavioral tasks provide the most informative signals. Third, we determined how much task data is required for reliable prediction. These analyses provide guidance for task selection and data collection during presurgical mapping. In addition, we examined how evaluation at the level of stimulation pairs influences interpretation of model performance, given the pair-based nature of electrical stimulation mapping.

## 2 Methods

### 2.1 Standard Protocol Approvals, Registrations, and Patient Consents

The study was approved by the Institutional Review Board of NYU Grossman School of Medicine. Written informed consent was obtained from all participants.

### 2.2 ECoG Recording and Preprocessing

#### Participants, tasks, and signal acquisition

Fourteen native English-speaking individuals with drug-refractory epilepsy undergoing intracranial monitoring at NYU Langone Health participated in this observational study. Subdural electrocorticography (ECoG) recordings were obtained using clinically implanted grid and strip electrodes for localization of the seizure onset zone. Coverage varied across participants and primarily included frontal, temporal, and perirolandic regions involved in speech and language. Demographic, implant, and electrode characteristics are summarized in Table 1.

**Table 1:** Patient, implant, and electrode cohort characteristics.

| Characteristic | Value |
| --- | --- |
| Total patients, n | 14 |
| Age, mean ± SD (range), years | 27.9 ± 12.7 (14.0–55.9) |
| Female sex, n (%) | 9 (64.3%) |
| Handedness, n | Right: 8; Left: 5; Ambidextrous: 1 |
| Implant hemisphere, n | Left: 12; Right: 0; Bilateral: 2 |
| Electrodes analyzed per patient, mean ± SD (range) | 93.9 ± 21.6 (48–127) |
| Total electrodes analyzed, n | 1314 |
| Electrodes tested with ESM, n | 586 |
| Critical electrodes identified by ESM, n (%) | 249 / 586 (42.5%) |
| Speech arrest electrodes, n (%) | 100 / 586 (17.1%) |
| Motor electrodes, n (%) | 93 / 586 (15.9%) |
| Language electrodes, n (%) | 56 / 586 (9.6%) |
| Unique cortical regions sampled, n | 24 |

Neural signals were recorded while participants performed a battery of speech production tasks separate from electrical stimulation mapping. Tasks were designed to overlap with clinical stimulation mapping procedures and include the same items tested during stimulation [20, 21]. The speech production tasks included:

#### Auditory Word Repetition (AR)

Repeating heard words.

#### Auditory Word Naming (AN)

Naming a word based on an auditory definition.

#### Sentence Completion (SC)

Completing the final word of an auditory sentence.

#### Visual Word Reading (VR)

Reading aloud visually presented words.

#### Picture Naming (PN)

Naming a word based on a colored line drawing.

Each participant completed the same 50 target words, presented once in AN and SC and twice in the remaining tasks, yielding 400 trials per participant. Electrodes with persistent artifacts or poor signal quality were excluded.

Signals were re-referenced using a common average reference (CAR), in which the mean across all electrodes was subtracted at each time point to reduce spatially diffuse noise. Recordings were segmented into epochs spanning -250 to 500 ms relative to speech onset for consistent temporal alignment across trials. Neural activity was baseline normalized using a 500-ms pre-event interval preceding stimulus onset, and signals were converted to z-scores based on this baseline.

Primary analyses focused on high-gamma activity (70–150 Hz), which is closely associated with local cortical activation and is widely used in electrocorticography studies of speech and functional mapping [5, 7, 9, 20, 21]. High-gamma analytic amplitude was obtained by bandpass filtering CAR-referenced signals (70–150 Hz), applying the Hilbert transform, and extracting the analytic amplitude envelope.

### 2.3 Ground-Truth Labels from Electrical Stimulation Mapping

Ground-truth labels were defined using clinical electrical stimulation mapping (ESM), performed independently of experimental recordings, as described in [22]. During ESM, bipolar stimulation was delivered between electrode pairs while participants performed standard mapping tasks. Both electrodes were labeled critical if stimulation reliably elicited a behavioral deficit and non-critical if no deficit was observed. Electrodes not tested were excluded.

#### Functional Label Definitions

Functional labels were defined based on the observed behavioral response during stimulation. Motor-positive electrodes were those in which stimulation produced reproducible motor deficits. Speech arrest was assessed in electrodes without motor responses and was defined as transient inability to speak despite preserved motor function [23]. Language-positive electrodes were those in which stimulation produced higher-order language deficits. Under the combined function definition, electrodes were labeled positive if stimulation produced any of the three deficits (motor, speech arrest, or language), reflecting a broader clinical definition of eloquent cortex. Behavioral criteria for each category are summarized in Table 2.

**Table 2:** Behavioral criteria used to define functional labels.

| Category | Behavioral criteria |
| --- | --- |
| Motor | Reproducible motor responses elicited by stimulation, including movements of the tongue, face, limbs, eyes, head, or shoulders, as well as articulatory motor responses (e.g., tongue, jaw, or laryngeal movement). |
| Speech arrest | Transient inability to speak during stimulation despite preserved motor function and absence of generalized motor impairment. |
| Language | Higher-order language disturbances, including naming impairment, word-finding difficulty, sentence completion errors, impaired comprehension, delayed responses, or paraphasic errors. |

### 2.4 Feature Extraction for Predicting Electrode Criticality

Feature extraction was performed independently for each electrode and trial to capture anatomical localization and functional network interactions. The primary representation included anatomical region encoding and graph-theoretic connectivity features, selected based on supplementary comparisons with alternative feature sets (Supplementary Section 3). Combining anatomical and connectivity features provided stable performance across domains, with statistically significant improvements for speech arrest prediction, compared to anatomical features alone (paired Wilcoxon signed-rank test, p<0.001), while more complex signal-derived features provided limited additional benefit.

#### 2.4.1 Anatomical region encoding

Each electrode was assigned to one of 24 cortical regions derived from the Desikan–Killiany–Tourville atlas [24]. Region identity was encoded using a one-hot vector representation, allowing the classifier to incorporate anatomical information and known functional specialization of cortical regions. Region lists and electrode distributions are provided in Supplementary Table 1, with cortical parcellation shown in Supplementary Figure 1.

#### 2.4.2 Graph-theoretic connectivity features

Functional interactions between electrodes were computed for each trial as absolute Pearson correlation between electrode time series. The resulting matrices were treated as weighted, undirected graphs, with electrodes as nodes and correlations as edge weights.

From each trial-specific network, we extracted three graph-theoretic measures for each electrode: strength, eigenvector centrality, and clustering coefficient. Strength reflects overall connectivity between an electrode and the rest of the network. Eigenvector centrality captures the influence of an electrode within the network by accounting for both its direct connections and the connectivity of its neighbors. The clustering coefficient quantifies the extent to which an electrode’s neighboring nodes are interconnected, reflecting local network organization. These measures capture complementary aspects of network organization, and are widely used in studies of brain connectivity [15, 17]. Additional details are provided in Supplementary Section 2.

Alternative connectivity representations, including Node2Vec embeddings and Graph Neural Network models [25, 26], were also evaluated but did not consistently improve performance (Supplementary Section 4).

### 2.5 Classification Framework

Because each electrode was sampled across multiple trials, classification was performed at the trial level with aggregation to obtain electrode-level predictions. For each trial, the classifier estimated the probability that the electrode was functionally critical based on anatomical and connectivity features. Trial-level probabilities were averaged to produce a single electrode-level probability:

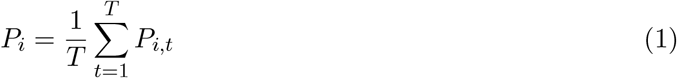

where *P*_*i,t*_ denotes the predicted probability for electrode *i* on trial *t*, and *T* is the number of trials. This aggregation strategy yielded more consistent performance than averaging neural signals prior to extracting the connectivity values (Supplementary Section 5).

We evaluated multiple machine learning classifiers, including linear and radial basis function support vector machine (SVM), logistic regression, random forest, decision tree, and multilayer perceptron models. The linear SVM provided the most consistent performance, with strong discrimination and stable threshold-dependent classification metrics. To account for class imbalance, balanced class weights were applied during training, increasing the penalty for misclassification of critical electrodes. Detailed comparisons of classifier performance are provided in Supplementary Section 6.

### 2.6 Evaluation Metrics

The classifier produces a probability estimate for each electrode. To obtain binary predictions, a decision threshold was applied, above which electrodes were classified as critical and below which they were classified as non-critical. Because performance depends on this threshold, both threshold-independent and threshold-dependent metrics were evaluated to capture complementary aspects of model behavior.

#### Threshold-independent metrics

Receiver operating characteristic area under the curve (ROC– AUC) measures the ability of the model to distinguish critical from non-critical electrodes across thresholds. It reflects the probability that a randomly selected critical electrode receives a higher predicted probability than a non-critical electrode.

Precision–recall area under the curve (PR–AUC) evaluates the trade-off between precision (positive predictive value) and recall (sensitivity) and is particularly informative in imbalanced datasets where critical electrodes are relatively uncommon. To account for differences in class prevalence across labeling definitions, we additionally report PR–AUC lift:

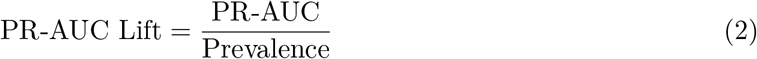

where Prevalence equals the percentage of critical (positive) samples among all samples. PR–AUC lift expresses performance relative to chance, with 1 indicating chance-level performance and higher values indicating above chance accuracy.

#### Threshold-dependent metrics and calibration

Binary classification performance under a chosen decision threshold was evaluated using F1 score and balanced accuracy:

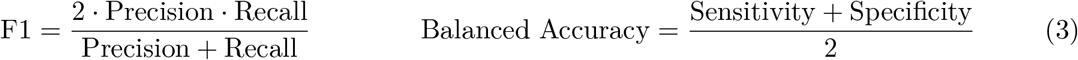

F1 score reflects the balance between sensitivity and precision, while balanced accuracy accounts for class imbalance by weighting sensitivity and specificity equally. Decision thresholds were selected by sweeping candidate thresholds to maximize F1 score across training subjects and then applied unchanged to the held-out subject.

Model performance was evaluated using leave-one-subject-out (LOSO) cross-validation to assess generalization across individuals. In each fold, models were trained on all but one subject and tested on the held-out subject, ensuring independence between training and evaluation data. Metrics were computed separately for each held-out subject and reported as mean ± standard deviation across subjects to reflect inter-subject variability and the consistency of model performance across individuals.

### 2.7 Data and Code Availability

The analysis framework and source code supporting this study is publicly available at https://github.com/nikaemami/ecog-eloquent-cortex-prediction. De-identified data analyzed in this study may be available from the corresponding author upon reasonable request, subject to institutional review board approval and data use agreements to protect patient privacy.

## 3 Results

### 3.1 Effect of Criticality Definition on Prediction Performance

Electrical Stimulation Mapping (ESM) identifies electrodes that are critical across functional domains (motor, speech arrest, or language). We suspect that the relationship between electrophysiological features and labels is function-dependent. Therefore, we develop three separate classifiers for the three functions. Each classifier predicts the ground truth ESM positive labels based on the electrophysiological features. Similarly, we develop a single classifier using combined labels to identify electrodes critical for any of the three functions. Lastly, we develop a *combined max-fusion* classifier, which applies all three single function classifiers and uses the highest predicted probability. Table 3 summarizes cross-validated performance. Receiver operating characteristic (ROC) and precision–recall (PR) curves across leave-one-subject-out folds are provided in Supplementary Section 7. Results are obtained with data from all tasks and the combined region + connectivity feature set.

**Table 3:** Model performance by criticality definition (mean ± SD across subjects).

| Criticality | ROC–AUC | PR–AUC | Bal Acc | F1 | Prev | PR–AUC Lift |
| --- | --- | --- | --- | --- | --- | --- |
| Motor | $0.929 \pm 0.061$ | $0.755 \pm 0.191$ | $0.872 \pm 0.084$ | $0.753 \pm 0.143$ | 0.165 | 4.561 |
| Speech Arrest | $0.793 \pm 0.103$ | $0.550 \pm 0.196$ | $0.742 \pm 0.125$ | $0.536 \pm 0.233$ | 0.198 | 2.772 |
| Language | $0.761 \pm 0.160$ | $0.385 \pm 0.167$ | $0.675 \pm 0.149$ | $0.284 \pm 0.189$ | 0.109 | 3.519 |
| Combined (Single Model) | $0.825 \pm 0.075$ | $0.739 \pm 0.150$ | $0.704 \pm 0.113$ | $0.612 \pm 0.238$ | 0.421 | 1.754 |
| Combined (Max-Fusion) | $0.848 \pm 0.091$ | $0.758 \pm 0.167$ | $0.732 \pm 0.126$ | $0.686 \pm 0.239$ | 0.421 | 1.800 |

*Abbreviations:* ROC–AUC = area under the receiver operating characteristic curve; PR–AUC = area under the precision–recall curve; F1 = F1 score; Bal Acc = balanced accuracy; Prev = prevalence of positive labels; PR–AUC Lift = PR–AUC divided by prevalence.

Model performance differed across criticality definitions. Motor-only labels showed the strongest discrimination (ROC–AUC 0.929 ± 0.061, PR-AUC 0.755 ± 0.191), suggesting relatively consistent electrophysiologic signatures across patients. Speech arrest–only labels also performed well (ROC– AUC 0.793 ± 0.103, PR-AUC 0.550 ± 0.196), with detection rates nearly three times above chance. Language-only labels were more challenging to predict (ROC–AUC 0.761 ± 0.160, PR-AUC 0.385 ± 0.167), consistent with the broader and more distributed organization of higher-order language functions.

For combined function prediction, both classifiers performed below motor but above speech arrest and language. The combined max-fusion strategy showed improved discrimination compared with the single model trained on combined function labels (ROC–AUC 0.848 vs 0.825; PR–AUC 0.758 vs 0.739), and also improved decision-level performance (F1 score 0.686 vs 0.612; balanced accuracy 0.732 vs 0.704). This indicates that while both approaches capture similar underlying electrophysiologic information, the max-fusion approach improves classification when identifying electrodes critical for any functional deficit. However, enrichment over chance decreased for combined function labels (PR–AUC lift 1.754 and 1.800 vs 2.772–4.561 for individual criticality), consistent with the higher prevalence of positive electrodes (0.421 vs 0.109–0.198), which reduces relative enrichment despite increased overall performance.

Additional threshold-dependent metrics, including positive predictive value (PPV) and negative predictive value (NPV), are summarized in Supplementary Table 8. Across subjects, NPV remained high for all models (0.849 to 0.964), indicating that electrodes predicted as non-critical were typically confirmed by stimulation testing. The combined max-fusion approach showed the highest sensitivity (0.844) but specificity was lower relative to individual models, reflecting increased tendency to classify electrodes as critical. In contrast, PPV, reflecting the proportion of predicted critical electrodes confirmed by stimulation, was lower and more variable across models (0.215 to 0.741), consistent with class imbalance and increased positive predictions.

Taken together, predictive performance depended strongly on how functional criticality was defined. More focal functions, particularly motor and speech arrest, showed more spatially coherent and electrophysiologically distinct patterns, leading to higher prediction accuracy. In contrast, combined labeling improved overall detection of critical sites but introduced greater heterogeneity and reduced functional specificity.

#### Visualization of Results

To further examine spatial organization, we visualized projections of ground-truth ESM labels and model-predicted probability maps for each criticality definition using the Mithra visualization toolbox [27] (Figure 1).

**Figure 1:**
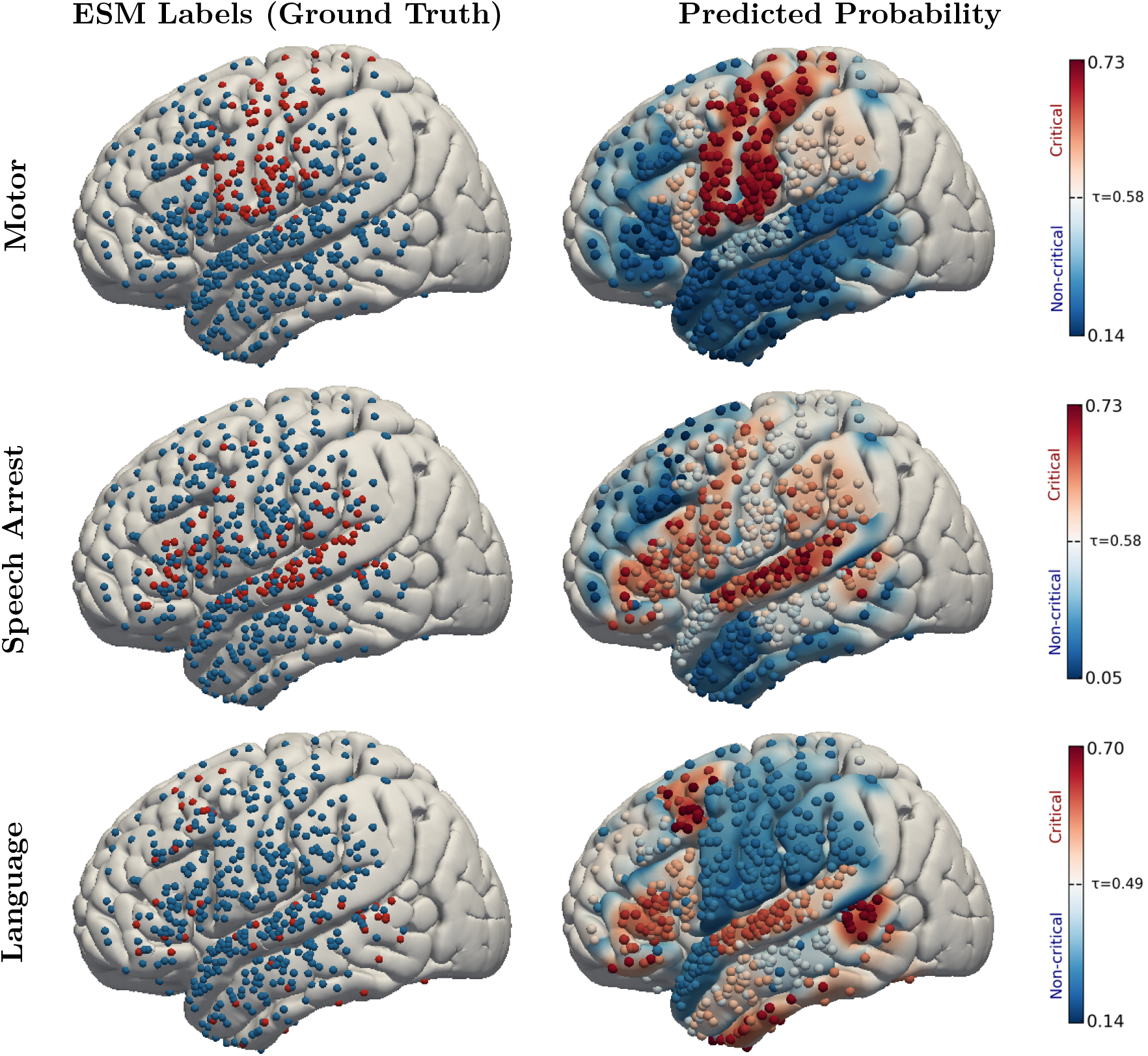
Across-subject projections of ESM-positive electrodes (left) and model-predicted probability of criticality (right) for motor, speech arrest, and language (*N* = 14 subjects).

Motor-critical sites exhibited a highly focal distribution centered over the central sulcus, with predicted probability maps closely mirroring ground-truth localization. Speech arrest sites showed a coherent pattern broadly tracking along perisylvian regions, while language-only labels were more spatially dispersed across frontal and temporal cortex, and predicted probability maps showed more diffuse activation.

### 3.2 Effect of Task Composition

We next examined how behavioral task selection influences prediction of functionally critical cortex. Because different language tasks engage partially overlapping but distinct neural systems, certain tasks may provide more informative signals for identifying specific functional deficits. To evaluate this, we trained and tested models using ECoG recordings from individual tasks, all pairwise and three-task combinations, and the full task set. Analyses were performed separately for motor, speech arrest, and language labels while holding all other aspects of the modeling framework constant, allowing isolation of the effect of task type on the predictive performance. Each task was evaluated for its ability to predict all functional domains. Model performance was evaluated using ROC–AUC and PR–AUC lift, and task configurations were ranked based on their average. PR–AUC lift was used because the prevalence of stimulation-positive labels differed across task configurations, as not all participants completed every task, leading to variation in sample size and class balance. Overall, task-related differences were modest, with certain tasks showing slightly stronger performance depending on the functional domain. Results are shown in Figure 2, with full results in Supplementary Section 8.

**Figure 2:**
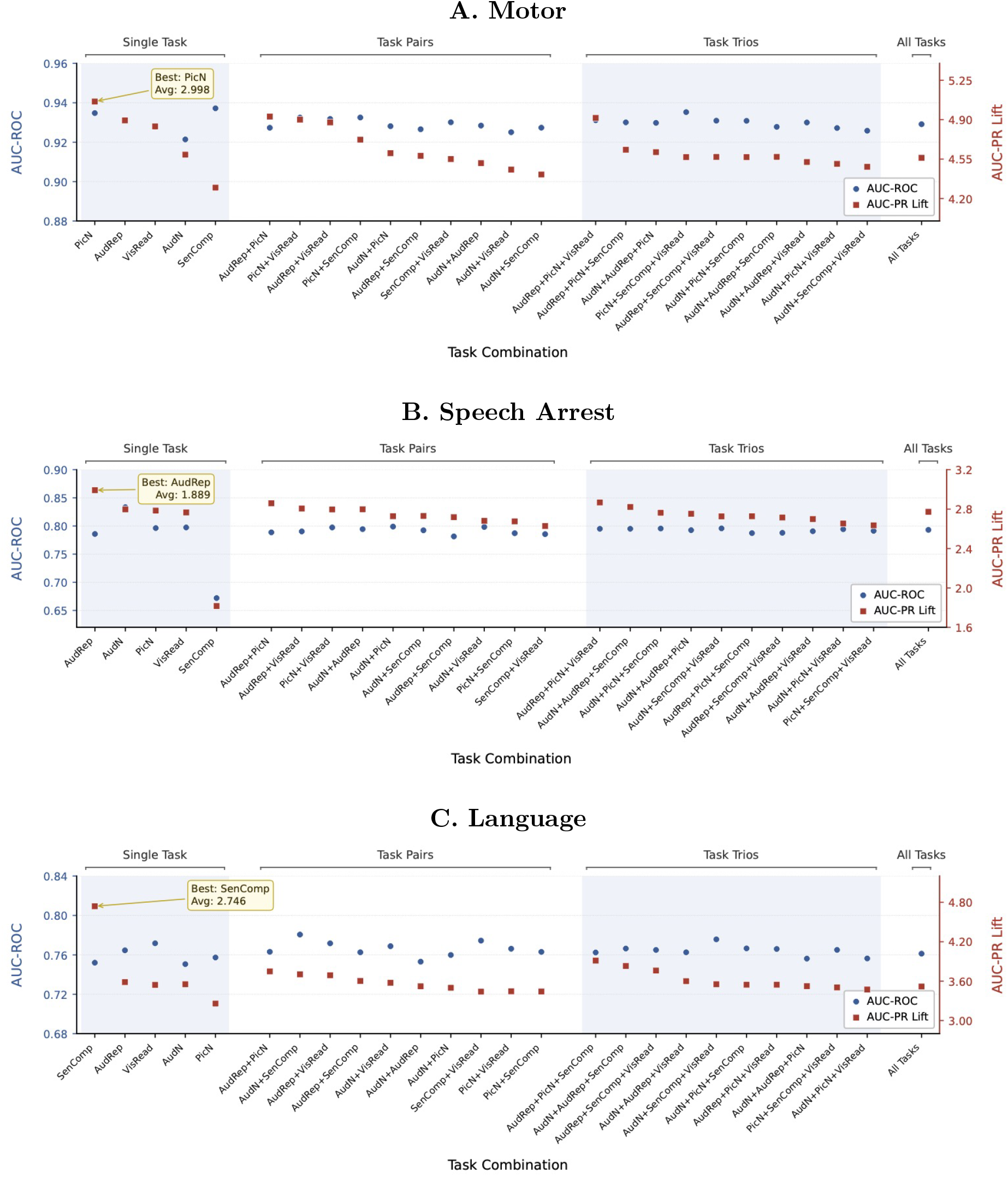
Model performance across task configurations for predicting motor, speech arrest, and language critical cortex. Each point represents a task configuration (single tasks, pairs, trios, or all tasks), with circles indicating ROC–AUC and squares indicating PR–AUC lift.

#### Motor

Motor-critical electrodes were detected reliably across nearly all task configurations. Among individual tasks, picture naming yielded the strongest performance (ROC–AUC 0.935 ± 0.055, PR– AUC 0.775 ± 0.175, lift 5.061), closely followed by auditory repetition and visual reading. Task combinations maintained similarly high performance but did not substantially exceed the best single-task models. Models incorporating all tasks performed comparably (ROC–AUC 0.929 ± 0.061, PR–AUC 0.755 ± 0.191, lift 4.561), indicating that motor-critical cortex can be identified robustly across behavioral conditions. This relatively consistent performance suggests that motor-critical electrodes are associated with stable, reproducible features that generalize across paradigms.

#### Speech Arrest

For speech arrest, performance showed modest variation across tasks, suggesting that some tasks may provide more informative signals than others. Among single-task models, auditory repetition yielded the strongest performance (ROC–AUC 0.786 ± 0.096, PR–AUC 0.536 ± 0.198, PR–AUC lift 2.992), followed by auditory naming and picture naming. Combining tasks provided modest improvements in some cases but did not consistently outperform the best individual task. Several task pairs and trios achieved performance comparable to auditory repetition alone, but none consistently exceeded it. Incorporating all five tasks produced similar performance (ROC–AUC 0.793±0.103, PR–AUC 0.550±0.196, lift 2.772), indicating that additional tasks did not substantially improve detection of speech arrest–critical cortex beyond the most informative paradigms.

#### Language

Language-critical sites showed greater variability across tasks. Among individual tasks, sentence completion showed the highest performance (ROC–AUC 0.752 ± 0.175, PR–AUC 0.511 ± 0.165, lift 4.741), outperforming tasks such as picture naming and auditory naming. Combining tasks resulted in modest improvements relative to weaker individual tasks but did not consistently exceed the best single-task model. Models incorporating all tasks performed similarly to many task combinations (ROC–AUC 0.761 ± 0.160, PR–AUC 0.385 ± 0.167, lift 3.519), suggesting that adding additional paradigms does not consistently enhance detection of language-critical cortex beyond the most informative tasks.

### 3.3 Effect of Number of Trials

In the preceding analysis, auditory repetition and picture naming emerged among the most informative tasks for predicting functionally critical cortex across domains. We therefore examined how predictive performance depends on the number of available trials within these tasks, an important practical consideration for presurgical mapping.

Each task included 100 trials per participant. To evaluate the impact of trial quantity, we varied the number of trials per electrode (1, 5, 10, 25, 50, 75, and 100), randomly sampling trials without replacement and averaging predicted probabilities across five subsamples to obtain stable estimates. Statistical comparisons were performed using bootstrap resampling of electrodes (5,000 iterations), with 95% confidence intervals computed for differences in ROC–AUC and PR–AUC between trial counts. This analysis was performed separately for motor, speech arrest, and language labels while holding the modeling framework constant. Overall, predictive performance improved from 1 to approximately 10 trials, with limited additional benefit beyond this range. This pattern was most pronounced and statistically reliable for speech arrest. Results are summarized in Figure 3, with full numerical results in Supplementary Section 9.

**Figure 3:**
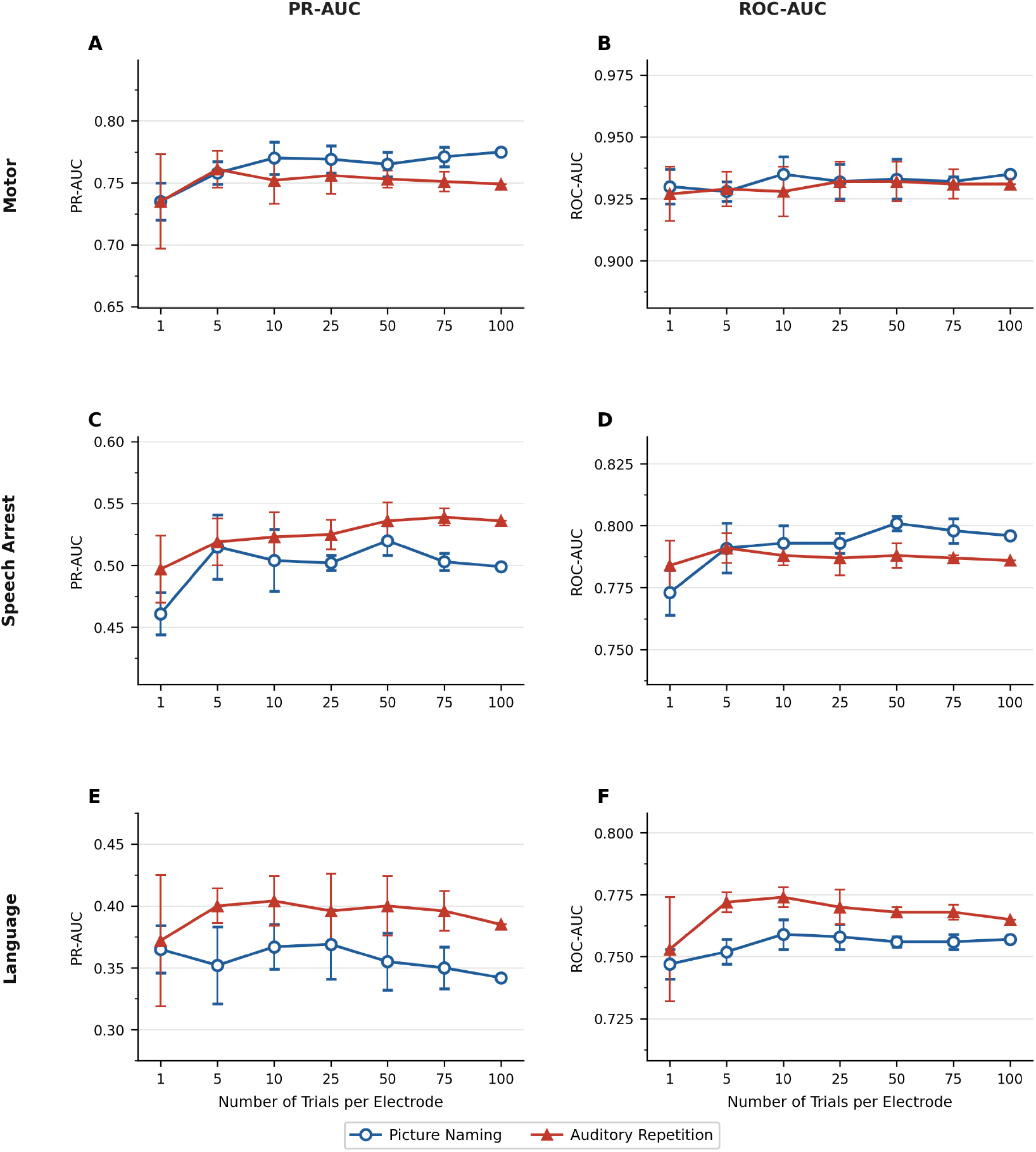
Effect of trial count on predictive performance for picture naming (blue circles) and auditory repetition (red triangles). Rows correspond to speech arrest, motor, and language critical labels. Left column shows PR–AUC and right column shows ROC–AUC. Error bars represent standard deviation across five random subsamples.

#### Motor

Motor-critical electrodes were identified reliably even with minimal sampling. Using a single trial yielded strong discrimination for both picture naming (ROC–AUC 0.930 ± 0.007, PR–AUC 0.735 ± 0.015) and auditory repetition (ROC–AUC 0.927 ± 0.011, PR–AUC 0.735 ± 0.038). Performance showed small numerical increases with additional trials (picture naming PR–AUC 0.770 at 10 trials and 0.775 with all trials), but remained high across trial counts, with no statistically significant differences. These findings suggest that motor-critical cortex exhibits stable, readily detectable features that do not require large numbers of trials.

#### Speech Arrest

Predictive performance improved from 1 to approximately 10 trials and plateaued thereafter. Using a single trial per electrode yielded moderate discrimination for both picture naming (ROC–AUC 0.773 ± 0.009, PR–AUC 0.461 ± 0.017) and auditory repetition (ROC–AUC 0.784 ± 0.010, PR–AUC 0.497 ± 0.027). Performance increased with additional trials, with gains observed up to approximately 10 trials for both tasks. Statistical analysis supported improvement between 1 and 10 trials for both ROC–AUC and PR–AUC. Beyond 10 trials, gains were small, and no significant improvement was observed between 10 and 100 trials. Small fluctuations at higher trial counts were consistent with variability rather than systematic changes.

#### Language

Predictive performance for language-critical cortex showed modest variation across trial counts, with a slight increase from 1 to approximately 10 trials for both tasks (ROC–AUC increased from approximately 0.75 to 0.77 between 1 and 10 trials for both auditory repetition and picture naming). PR–AUC showed similar but more variable behavior without a consistent increasing trend. However, these differences did not reach statistical significance (bootstrap 95% CI), consistent with higher variability and lower statistical power at low trial counts. Beyond 10 trials, performance remained stable across both tasks, with no consistent improvement up to 100 trials. Overall performance remained lower than in motor and speech arrest domains.

### 3.4 Effect of Evaluation Level: Electrode versus Stimulation Pair

Electrical stimulation mapping (ESM) is performed between electrode pairs, whereas the preceding analyses evaluated model performance at the level of individual electrodes. To examine how this difference affects interpretation, we performed an additional evaluation at the level of stimulation pairs. For this analysis, ground truth was defined from raw ESM outcomes, with pairs labeled positive if stimulation produced a functional deficit and negative otherwise. A stimulation pair was classified as positive if either electrode in the pair was predicted as critical, using the same decision threshold as in the electrode-level analysis.

Across all functional domains, pair-level evaluation resulted in higher sensitivity compared to electrode-level evaluation (motor: 0.924 vs 0.817; speech arrest: 0.761 vs 0.682; language: 0.773 vs 0.672; Table 4). This came at the cost of lower specificity (motor: 0.880 vs 0.926; speech arrest: 0.696 vs 0.802; language: 0.551 vs 0.679), indicating that pairs without deficits were more often classified as positive. Overall performance (ROC–AUC and PR–AUC) showed only modest differences between evaluation levels. These findings suggest that the model often identifies at least one electrode within stimulation pairs that produce deficits, even when exact agreement at the individual electrode level is limited. This pattern is consistent with the spatial uncertainty inherent in bipolar stimulation, wherein the functional effect is observed for a pair rather than a single electrode.

**Table 4:** Change in performance (stimulation-pair minus electrode-level; mean across subjects).

| Domain | ΔSensitivity | ΔSpecificity | ΔROC-AUC | ΔPR-AUC |
| --- | --- | --- | --- | --- |
| Motor | +0.107 | -0.046 | +0.003 | +0.014 |
| Speech Arrest | +0.079 | -0.106 | -0.019 | -0.027 |
| Language | +0.101 | -0.128 | -0.024 | -0.019 |

## 4 Discussion

Identifying cortical regions essential for speech and language is critical for preserving neurological function during brain surgery. Electrical stimulation mapping (ESM) remains the clinical gold standard but is time-consuming, limited by incomplete sampling, and constrained by patient tolerance and testing time [3, 4]. Early ECoG studies showed that high-gamma activity during task performance can identify functionally relevant cortex [5–7]. In this study, we build on prior approaches by incorporating graph-theoretic measures of functional connectivity from ECoG to identify stimulation-defined critical cortical sites. Predictive performance varied across functional domains, tasks, and trial counts, providing insight into the organization of eloquent cortex and practical considerations for presurgical mapping.

We observed marked differences in predictive performance across functional domains. Previous ECoG studies show that higher-order language function is supported by distributed, task-dependent networks rather than a single focal region [8, 10, 12]. Consistent with this, motor-critical electrodes were identified with the highest accuracy, followed by speech arrest, whereas language-critical cortex was more difficult to predict. This pattern likely reflects underlying neurophysiologic organization: motor representations are relatively focal and consistent across individuals, whereas higher-order language functions are more distributed and involve multiple interacting processes, resulting in greater variability across subjects. These findings align with network-based models of language organization [15, 16, 19] and with the broader, less spatially consistent distributions observed for language-critical electrodes. Our results extend these prior observations by demonstrating that such domain-specific differences persist in a cross-subject predictive framework, and suggest that network-level interactions can contribute to identifying functionally critical cortex.

Prior machine learning approaches have largely relied on local spectral ECoG features evaluated within individual subjects [14], without explicitly modeling interactions between cortical sites. Graph-theoretic measures derived from functional connectivity characterize such interactions. Network neuroscience shows that measures such as strength, eigenvector centrality, and clustering coefficient capture complementary aspects of brain organization [15–18]. Recent work further supports their relevance for identifying stimulation-defined functional sites, showing that language-critical electrodes act as connectors between functional subnetworks [19]. In our framework, combining anatomical and connectivity features resulted in a statistically significant improvement for speech arrest prediction, confirmed by a paired Wilcoxon signed-rank test on pooled electrodes across subjects (p<0.001), while motor prediction remained largely driven by anatomical localization and language prediction remained less consistent across subjects (Supplementary Section 3). More complex connectivity representations, including Node2Vec embeddings and Graph Neural Networks, did not consistently improve performance despite increased model complexity, supporting the use of interpretable graph-theoretic features in a practical presurgical setting (Supplementary Section 4).

In addition to domain-specific differences, the definition of functional criticality influenced model behavior. Combined labeling, which classifies electrodes as critical if stimulation produces any deficit, reflects common clinical practice [3, 4]. Among the two prediction strategies for this, combined max-fusion of the results from three separate classifiers achieved higher performance than a single model trained on combined labels. This approach increased sensitivity and improved detection of electrodes critical for any function, but reduced specificity. In contrast, domain-specific models preserved functional and anatomical specificity, with motor-specific models achieving the highest performance. Across models, negative predictive value remained consistently high, whereas positive predictive value was more variable. These results highlight an important tradeoff. Domain-specific models support more precise localization, while combined max-fusion improves sensitivity and broader identification of potentially critical regions across functions. Clinically, these findings suggest that combined max-fusion may be preferred when prioritizing sensitivity to avoid missing eloquent cortex, even at the cost of more false positives.

Behavioral task selection influenced predictive performance differently across domains. Prior studies show that different language paradigms engage partially distinct cortical regions [10, 12]. In our results, task selection had a more limited effect on predictive performance across subjects. Motor-critical electrodes were identified reliably across tasks, suggesting that distinguishing features generalize across behavioral conditions. This likely reflects the strong contribution of anatomical location, which remained highly informative even without task-specific information. In contrast, speech arrest and language prediction showed greater variability, with certain paradigms, particularly auditory repetition and picture naming for speech arrest, and sentence completion for language, providing more informative signals. Combining tasks did not consistently improve performance beyond the best individual paradigms. These results suggest that task selection can be optimized based on the functional domain of interest, although differences across tasks were modest relative to inter-subject variability.

The number of trials per electrode also showed domain-specific effects. Prior studies report variable sensitivity of ECoG for identifying stimulation-defined language sites across patients [9, 28], including analyses of practical considerations around task repetitions in mapping performance. However, the domain-specific effect of trial count on predictive performance has not been systematically examined. In our study, motor-critical electrodes were identified accurately even with minimal sampling, with stable performance across trial counts. Speech arrest prediction improved up to approximately 10 trials, with statistically significant gains, then plateaued. Language prediction showed greater variability and remained lower overall, suggesting that limitations are driven more by the discriminative power of the features rather than the number of trials. These findings indicate that reliable localization of some domains may be achievable with limited sampling, whereas improving prediction for more distributed functions such as language may require more informative features.

Evaluation at the level of electrode pairs provided additional insight. Because ESM is inherently pair-based, electrode-level evaluation may underestimate the model’s ability to identify functionally relevant cortex [11, 28]. Prior work shows that stimulation effects may reflect disruption of distributed cortical networks rather than isolated sites [3, 4], which can complicate precise localization. In our analysis, pair-level evaluation increased sensitivity, indicating that the model often assigns high probability to at least one electrode within stimulation pairs that produce deficits. Clinically, this may be desirable when avoiding resection of eloquent cortex, although reduced specificity suggests overestimation of spatial extent. These findings highlight the complementary roles of the two evaluation approaches: electrode-level analysis reflects spatial precision, whereas pair-level analysis aligns more closely with characteristics of ESM.

These findings have direct implications for presurgical functional mapping. Connectivity-based analysis of task-related ECoG activity may help prioritize electrodes for stimulation and identify regions at elevated functional risk, particularly when mapping is incomplete or limited by patient tolerance [3, 4]. These approaches are intended to complement, rather than replace, electrical stimulation mapping. They may help guide task selection, improve testing efficiency, and support clinical decision-making.

### Limitations

Electrode coverage was determined by clinical needs and therefore varied across participants, resulting in incomplete sampling of cortex. Ground-truth labels were based on stimulation mapping, which itself may be influenced by stimulation parameters and clinical interpretation. In addition, the present study focused on high-gamma activity and connectivity features derived from speech tasks, and other neural features or task paradigms may provide additional predictive information. Future studies in larger and more diverse cohorts will be important to assess generalizability and clinical applicability.

## Conclusion

Neural activity recorded during speech production tasks contains information predictive of stimulation-defined functional criticality. Graph-theoretic connectivity-based decoding provides a framework for extracting this information and offers insight into the electrophysiological organization of eloquent cortex. These results support the potential role of connectivity-informed analysis as a complementary tool to improve the efficiency and interpretation of presurgical functional mapping.

## 5 Acknowledgment

This work was supported by the National Science Foundation under Grant No. IIS-2309057 (Y.W., A.F.), and National Institute of Health R01NS109367, R01NS115929. A.M. is supported by NIH grant 1K23NS146693.

