## Supplementary Material for "Prediction of Stimulation-Defined Eloquent Cortex Using Graph-Theoretical Connectivity from Electrocorticography During Presurgical Mapping"

### Supplementary Materials

#### 1 Cortical region definitions and electrode distribution

Cortical regions were derived from the Desikan–Killiany–Tourville atlas [1] and grouped into 24 categories based on electrode coverage and anatomical relevance to speech and language processing. Table S1 summarizes the list of regions and the total number of electrodes assigned to each region across all subjects. A visualization of the cortical parcellation used in this study is provided in Figure S1.

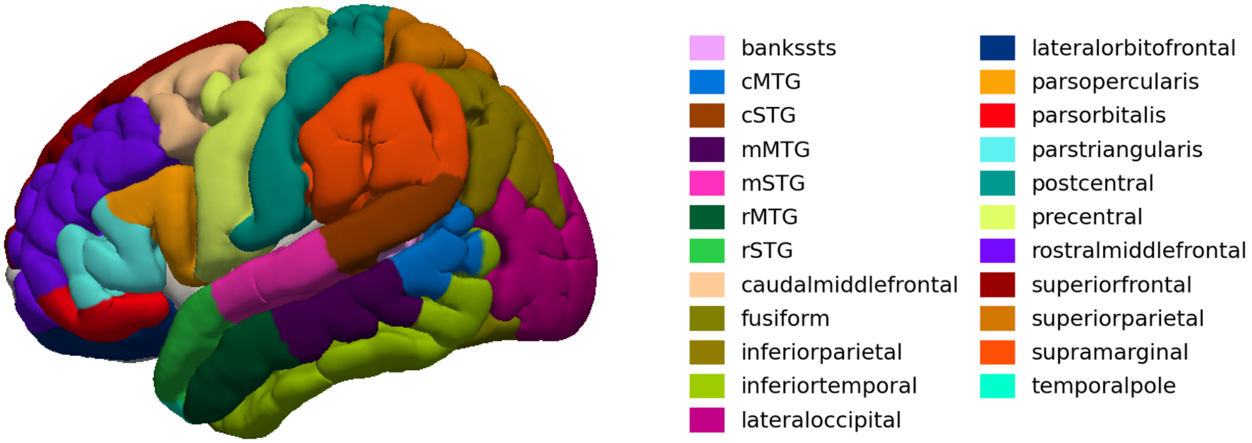


Figure S1: Cortical parcellation of the left hemisphere used in this study, shown on the FSL MNI152 pial surface. Each color corresponds to a distinct anatomical region from the Desikan-Killiany atlas.

Table S1: Electrode counts per cortical region across all subjects

| **Cortical region** | **Total electrodes, n** |
|---|---:|
| Caudal middle frontal | 20 |
| Lateral orbitofrontal | 3 |
| Pars opercularis | 26 |
| Pars orbitalis | 7 |
| Pars triangularis | 30 |
| Precentral | 49 |
| Rostral middle frontal | 26 |
| Superior frontal | 6 |
| Inferior parietal | 6 |
| Postcentral | 67 |
| Superior parietal | 2 |
| Supramarginal | 46 |
| cMTG | 19 |
| mMTG | 46 |
| rMTG | 42 |
| cSTG | 30 |
| mSTG | 48 |
| rSTG | 39 |
| Banks of STS | 7 |
| Fusiform | 11 |
| Inferior temporal | 32 |
| Temporal pole | 3 |
| Lateral occipital | 10 |
| Other | 11 |
| Total | 586 |

#### 2 Graph-theoretic connectivity feature definitions

**Functional connectivity** To quantify functional interactions between electrodes, we computed trial-specific connectivity matrices using the absolute Pearson correlation between electrode time series. For each trial, this yielded a symmetric weighted adjacency matrix of size $L \times L$, with $L$ denoting the number of electrodes:

$$W = \{w_{ij}\}, \quad w_{ij} = |\mathrm{corr}(x_i, x_j)| \tag{1}$$

where $x_i$ and $x_j$ denote neural signals recorded from electrodes $i$ and $j$.

These matrices were interpreted as weighted, undirected graphs, with electrodes as nodes and correlation values representing edge weights, enabling characterization of network topology using established graph-theoretic measures [2, 3].

From each trial-specific connectivity graph, we extracted three complementary measures for each electrode: strength, eigenvector centrality, and clustering coefficient, defined as below:

**Strength.** Strength quantifies the overall level of connectivity between an electrode and the rest of the network:

$$S(i) = \frac{1}{L-1} \sum_{j \neq i} w_{ij} \tag{2}$$

This measure reflects the average functional coupling of an electrode with other electrodes and its overall network integration.

**Eigenvector centrality.** Eigenvector centrality measures the relative influence of an electrode by accounting for both its direct connections and the connectivity of its neighbors. It is determined by the eigenvector $E = [E_i]$ of the adjacency matrix $W$ corresponding to the largest eigenvalue $\lambda$, satisfying

$$E_i = \frac{1}{\lambda} \sum_j w_{ij} E_j \tag{3}$$

Higher values indicate electrodes connected to other highly connected nodes, consistent with network hub structure.

**Clustering coefficient.** The clustering coefficient quantifies the degree to which an electrode's neighbors are themselves interconnected:

$$C_i = \frac{\sum_{j,k}(w_{ij}w_{jk}w_{ki})^{1/3}}{k_i(k_i - 1)} \tag{4}$$

where $k_i$ denotes the number of connections for electrode $i$. Because the connectivity matrix is not thresholded, the network is fully connected and $k_i = L - 1$ for all electrodes. This metric reflects local network organization and identifies electrodes embedded within tightly interconnected functional clusters.

### 3 Supplementary Analysis: Effect of Feature Set Composition

To evaluate the relative contributions of anatomical localization, functional connectivity, and signal-derived features, we compared multiple feature configurations using the same leave-one-subject-out framework and classifier settings. Performance metrics in this supplementary analysis were computed using pooled electrode-level predictions across all subjects to enable statistical comparisons between feature sets. All connectivity features were derived from the production-aligned window used in the main analyses.

Anatomical features were derived by assigning each electrode to one of cortical regions defined by the Desikan–Killiany–Tourville atlas [1], providing explicit spatial context to the classifier. Connectivity features were derived from trial-level Pearson correlation matrices and summarized using graph-theoretic metrics, including strength, eigenvector centrality, and clustering coefficient, as described in detail in the Methods section. These features capture complementary aspects of network integration and local connectivity [2, 3].

We also evaluated extended signal feature representations combined with anatomical and connectivity features. These included PCA-reduced signal features (capturing the main patterns in the time series) [4], mean and standard deviation (capturing overall activation magnitude and variability), and temporally downsampled signals at 10 Hz and 20 Hz (preserving coarse temporal dynamics while reducing dimensionality). These comparisons assess whether signal-level information provides complementary predictive value beyond anatomical and connectivity features.

Statistical comparisons between feature sets were performed using bootstrap resampling of pooled electrodes across all subjects (5,000 iterations) [5]. For each comparison, electrodes were resampled with replacement, and differences in performance metrics (ROC–AUC, PR–AUC, F1 score, and balanced accuracy) were computed across resamples. For threshold-dependent metrics (F1 score and balanced accuracy), each model was evaluated using its corresponding fixed global threshold determined during training. Statistical significance was determined based on whether 95% confidence intervals excluded zero, and bootstrap p-values were computed from the resampled difference distributions.

**Motor** Motor prediction performance was already strong using anatomical features alone (PR–AUC 0.56, ROC–AUC 0.90, F1 0.74, balanced accuracy 0.87; Figure S2). Connectivity features alone performed substantially worse, confirming that anatomical localization provides the dominant signal for identifying motor-critical cortex. Incorporating connectivity did not produce consistent improvements across evaluation metrics, and performance remained largely unchanged across feature sets. This pattern may reflect the relatively well-localized and anatomically consistent organization of motor cortex, which may already be adequately captured by anatomical features alone. These findings indicate that anatomical location captures the primary determinants of motor criticality.

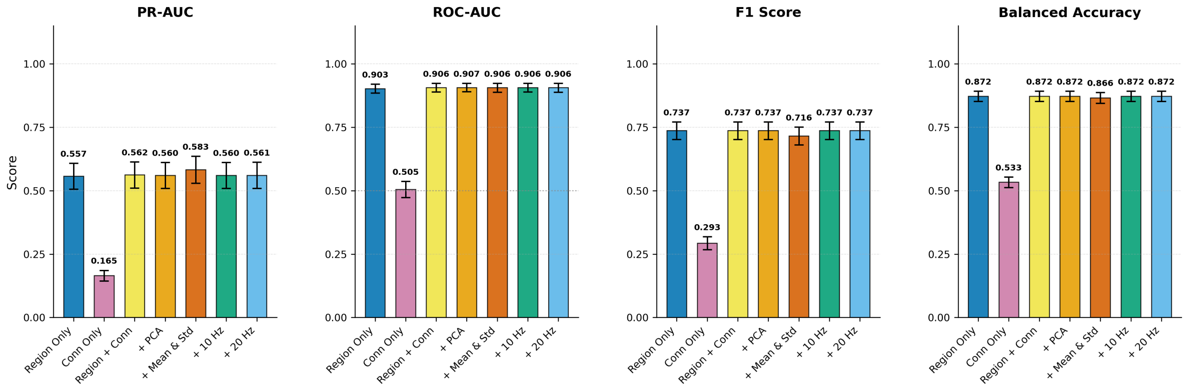


Figure S2: Effect of feature set composition on Motor prediction. Bars show pooled-electrode performance metrics across subjects; Error bars represent the bootstrap SE (5,000 resamples)

**Speech Arrest** For speech arrest prediction, anatomical features alone provided moderate discrimination (PR–AUC 0.38, ROC–AUC 0.76, F1 0.46, balanced accuracy 0.68; Figure S3). Connectivity features alone performed substantially worse across all metrics, indicating that network structure without anatomical context was insufficient for reliable classification.

Combining anatomical and connectivity features improved performance across all evaluation metrics, including ROC–AUC ($\Delta = +0.044$, 95% CI [0.018, 0.071], $p = 0.0004$), PR–AUC ($\Delta = +0.120$, 95% CI [0.051, 0.176], $p = 0.0002$), F1 score ($\Delta = +0.120$, 95% CI [0.068, 0.175], $p = 0.0002$), and balanced accuracy ($\Delta = +0.082$, 95% CI [0.039, 0.127], $p = 0.0012$). These differences were statistically significant based on bootstrap resampling of pooled electrodes across subjects, indicating that connectivity provides complementary information beyond anatomical localization. Extended signal-derived feature sets resulted in comparable performance to the combined anatomical and connectivity representation, suggesting limited additional benefit beyond this feature set.

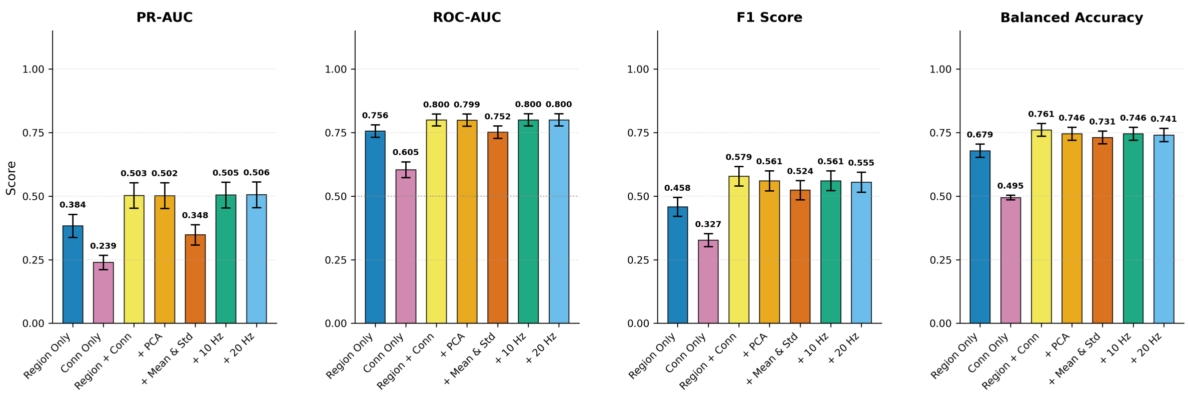


Figure S3: Effect of feature set composition on Speech Arrest prediction.

**Language** Language prediction showed only moderate performance using anatomical features alone (PR–AUC 0.23, ROC–AUC 0.75, F1 0.32, balanced accuracy 0.65; Figure S4), lower than that observed for motor and speech arrest predictions. Connectivity features alone performed poorly across all metrics, indicating limited predictive value in isolation. Incorporating connectivity and signal-derived features resulted in minimal change across evaluation metrics. Differences across feature sets were not consistently statistically significant across evaluation metrics. This may be influenced in part by the lower prevalence of language-critical electrodes in the dataset, which can increase variability and limit the ability to detect consistent improvements across feature sets. These findings suggest that language-critical cortex is more difficult to predict within this framework, and that neither anatomical location nor connectivity features provide consistently strong discriminative signatures. This may reflect the more distributed and heterogeneous functional organization of language cortex, resulting in weaker and more variable electrophysiologic predictors.

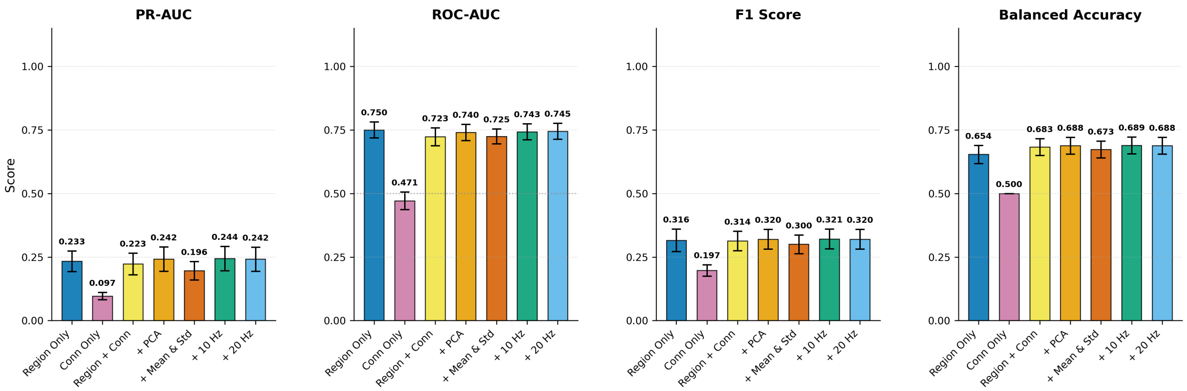


Figure S4: Effect of feature set composition on Language prediction.

**Summary** Across functional domains, anatomical localization emerged as the dominant predictor, while connectivity features alone were insufficient for reliable prediction. When combined with anatomical features, connectivity provided complementary information for speech arrest, with statistically significant improvements observed in pooled electrode-level analysis across metrics, but did not yield consistent improvements for motor or language prediction, suggesting that the utility of connectivity features may be domain-specific. These trends are further illustrated in Supplementary Figure S5 and the corresponding probability distribution (violin) plots (Figure S6), which show improved separation between critical and non-critical electrodes for speech arrest with connectivity features, but minimal change for motor and language.

Additional signal-derived features, including PCA-reduced representations, summary statistics, and temporally downsampled signals, did not provide consistent improvements beyond the combined anatomical and connectivity feature set. Based on these findings, we selected the combined Region + Connectivity feature representation derived from the production-aligned window for all subsequent analyses. This configuration provided strong performance for speech arrest and stable performance for motor and language predictions, while maintaining a simpler and physiologically interpretable feature representation compared to more complex features.

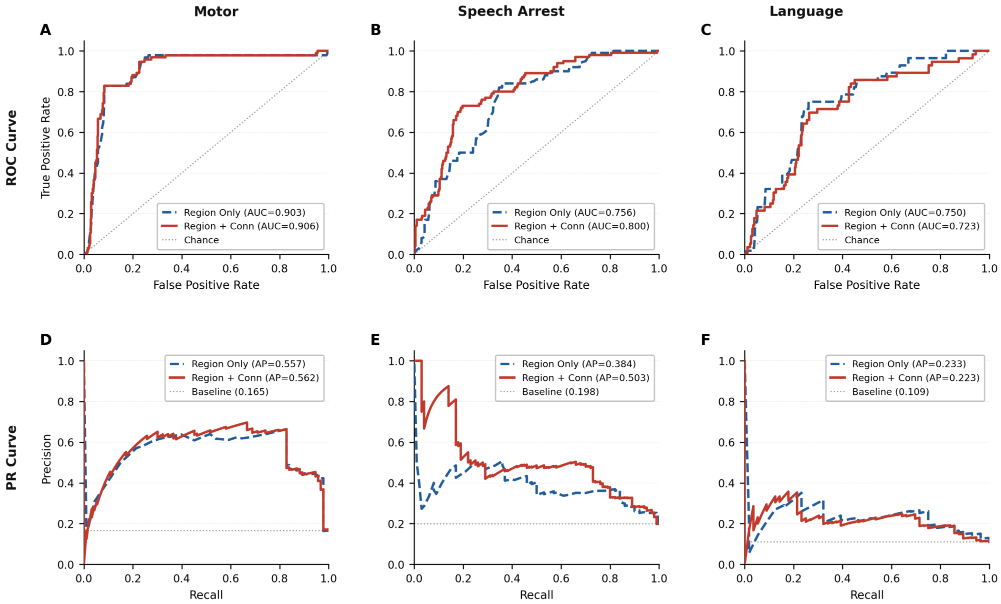


Figure S5: ROC curves (top row) and PR curves (bottom row) for models using anatomical features alone and combined anatomical and connectivity features, shown for motor (A,D), speech arrest (B,E), and language (C,F) prediction. Curves are computed using electrode-level predictions across subjects.

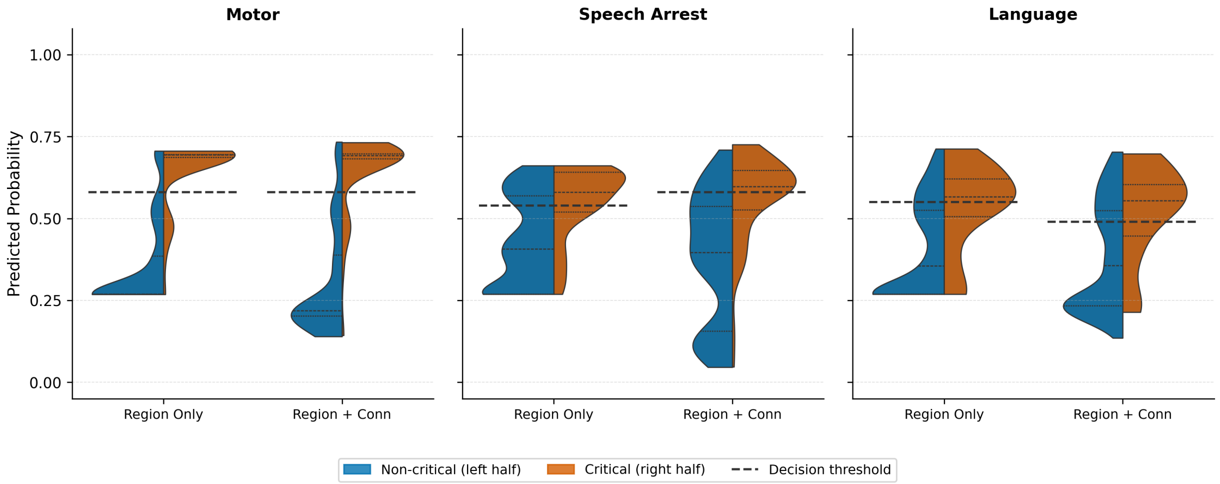


Figure S6: Predicted probability distributions for models using anatomical features alone and combined anatomical and connectivity features, shown for motor, speech arrest, and language. Each violin is split to show non-critical (left half) and critical (right half) electrodes, with dashed lines indicating decision thresholds.

To better align with the stimulation-based ground truth, we repeated the feature set comparison using pair-level predictions. As described in the main text (Section 3.4), pairs were labeled based on raw ESM outcomes, and predictions were defined at the pair level using the maximum probability of the two electrodes. Performance metrics were computed using pooled pair-level predictions across subjects. The results are shown below.

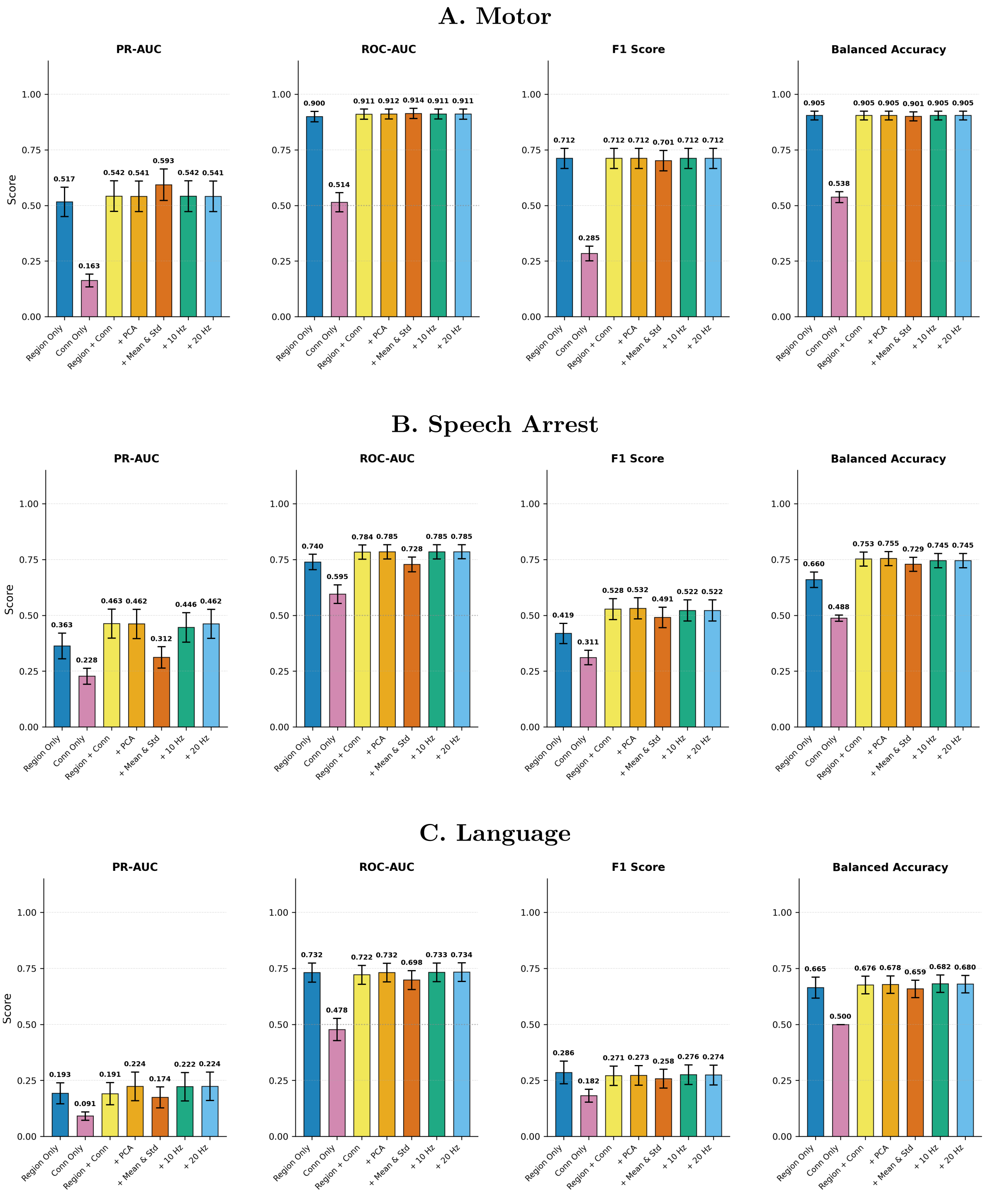


Figure S7: Effect of feature set composition using pair-level evaluation, shown for motor (A), speech arrest (B), and language (C) prediction. Bars show pooled pair-level performance metrics across subjects; error bars represent the bootstrap SE (5,000 resamples).

### 4 Supplementary Analysis: Effect of Connectivity Representation and Modeling Approach

To evaluate how different modeling strategies leverage inter-electrode connectivity for critical region prediction, we compared three classes of approaches: (1) classical graph-theoretic connectivity features combined with a static classifier, (2) Node2Vec graph embeddings combined with a static classifier, and (3) a graph neural network (GraphSAGE) operating directly on the connectivity graph.

Across all approaches, functional connectivity was estimated using absolute Pearson correlation between all electrode pairs within each trial, yielding symmetric undirected connectivity matrices. All models were evaluated using the identical leave-one-subject-out framework and production-aligned analysis window used throughout the main analyses, ensuring that differences reflect modeling strategy rather than data selection.

##### Classical Graph-Theoretic Features

In the first approach, connectivity graphs were summarized using established network metrics and provided as electrode-level features to a linear support vector machine classifier. Specifically, we extracted strength, eigenvector centrality, and clustering coefficient for each electrode, as defined in the Methods section. These features provide interpretable summaries of network topology while reducing the full connectivity matrix to compact, low-dimensional representations suitable for conventional classifiers [2, 3].

##### Node2Vec Embeddings

To obtain data-driven connectivity representations without imposing predefined summary statistics, we used Node2Vec to learn low-dimensional embeddings from trial-specific connectivity graphs [6].

**Graph construction.** For each subject–task–trial, electrodes were treated as nodes and edge weights were defined by the absolute Pearson correlation between electrode time series. Graphs were sparsified by retaining the top 20% strongest connections and ensuring overall connectivity.

**Embedding learning.** Node2Vec embeddings were learned independently for each trial using weighted random walks and a skip-gram objective [6]. We used 64-dimensional embeddings to capture electrode connectivity structure.

**Downstream classification.** Trial-level embeddings were combined with anatomical region encoding and evaluated using the same linear support vector machine leave-one-subject-out framework described in the Methods.

#### Graph Neural Network (GraphSAGE)

We evaluated a graph neural network approach using GraphSAGE [7], which directly operates on connectivity graphs and learns electrode representations through neighborhood aggregation.

Each trial was represented as a graph, with electrodes as nodes in a node-level classification task. Node features were derived from neural signals, while anatomical region encoding was added at the classifier stage. Functional connectivity graphs were constructed using absolute Pearson correlation between electrode signals, retaining the top 20% strongest connections while ensuring overall graph connectivity. GraphSAGE models were implemented using a three-layer architecture and evaluated with both mean and pooling neighborhood aggregation strategies, where node representations are updated by aggregating features from neighboring electrodes either through simple averaging (mean) or via a learned nonlinear pooling operation (max-pooling).

Learned node embeddings were combined with anatomical features and passed to a linear classifier to predict electrode criticality. Models were trained using weighted cross-entropy loss to address class imbalance and evaluated using the same leave-one-subject-out protocol as static classifiers. Only electrodes with definitive ESM labels contributed to the loss; untested electrodes were retained within the graph but masked during optimization. Trial-level predictions for each electrode were aggregated to produce electrode-level probabilities using the same procedure described in the Methods. Performance comparisons across connectivity modeling approaches are summarized in Figures S8–S10.

**Motor** Motor prediction showed strong performance across all connectivity representations. Classical connectivity features achieved slightly higher PR–AUC ($0.755 \pm 0.191$) compared with Node2Vec, while ROC–AUC, F1 score, and balanced accuracy were similar between the two approaches, indicating broadly comparable predictive performance overall. Graph neural network approaches produced slightly higher discrimination, with max-pooling aggregation achieving PR–AUC $0.798 \pm 0.178$ and ROC–AUC $0.943 \pm 0.062$. However, decision-level performance remained similar or slightly lower (F1 $0.742 \pm 0.143$, balanced accuracy $0.862 \pm 0.080$), indicating limited improvement in practical electrode-level classification.

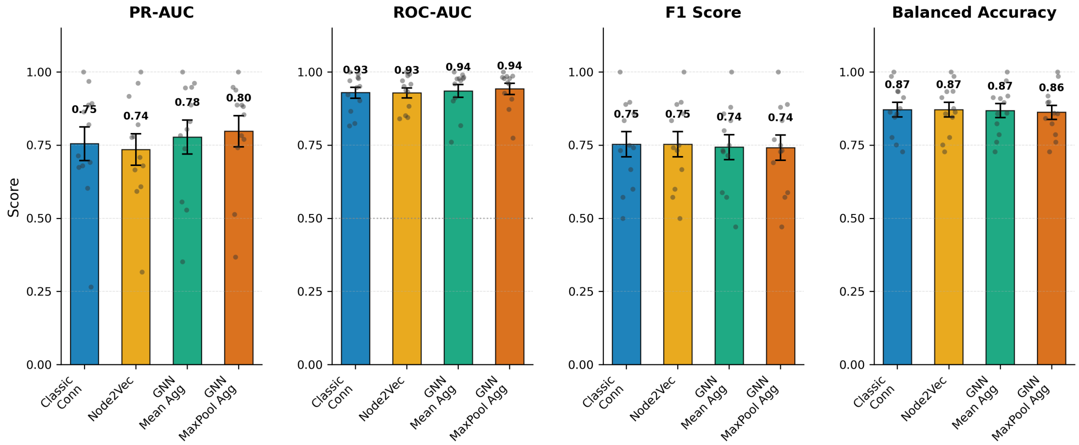


Figure S8: Connectivity representation and modeling approach for Motor prediction. Bars show mean ± SEM across LOSO folds; gray dots indicate individual fold values.

**Speech Arrest** For speech arrest prediction, classical graph-theoretic connectivity features achieved strong and balanced performance across all metrics (PR–AUC $0.550 \pm 0.196$, ROC–AUC $0.793 \pm 0.103$, F1 $0.536 \pm 0.233$, balanced accuracy $0.742 \pm 0.125$). Node2Vec embeddings yielded slightly higher discrimination (PR–AUC $0.569 \pm 0.190$, ROC–AUC $0.798 \pm 0.101$) but lower decision-level performance (F1 $0.521 \pm 0.184$, balanced accuracy $0.737 \pm 0.111$), indicating comparable discriminative capacity but less stable decision boundaries. Graph neural network approaches showed lower performance across both discrimination and decision-level metrics, indicating reduced reliability relative to static connectivity feature representations.

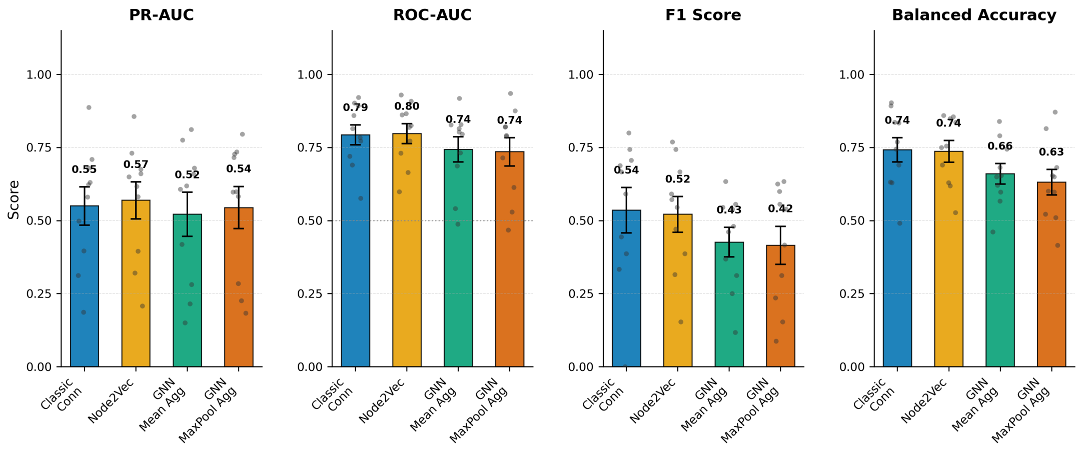


Figure S9: Connectivity representation and modeling approach for Speech Arrest prediction.

**Language** Language prediction showed modest variation across connectivity representations. Node2Vec achieved slightly higher performance than classical connectivity features across all evaluation metrics (PR–AUC 0.432±0.243 vs. 0.385±0.167, ROC–AUC 0.798±0.158 vs. 0.761±0.160, F1 $0.314 \pm 0.204$ vs. $0.284 \pm 0.189$, balanced accuracy $0.705 \pm 0.154$ vs. $0.675 \pm 0.149$). Graph neural network approaches did not provide improvement over classical connectivity features, with lower or comparable performance across all metrics.

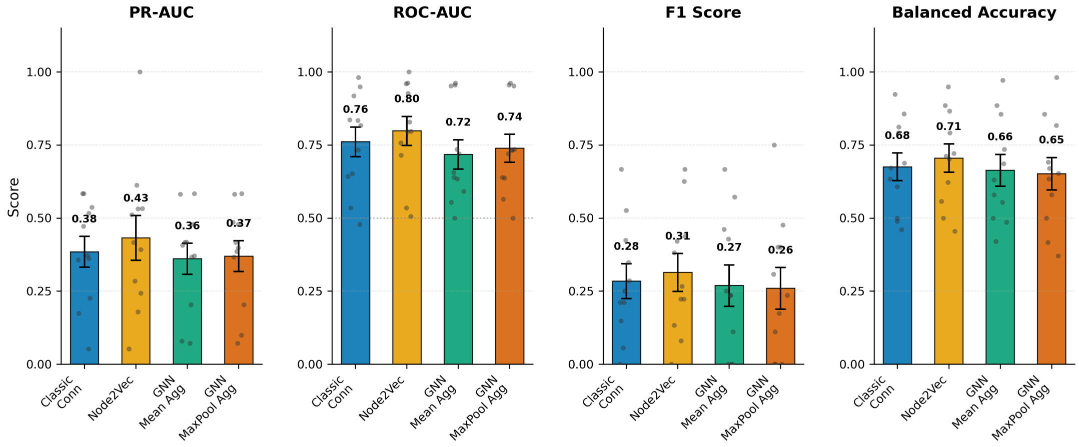


Figure S10: Connectivity representation and modeling approach for Language prediction.

**Summary** Across functional domains, classical graph-theoretic connectivity features provided strong and consistent performance across all metrics. While Node2Vec achieved comparable or slightly higher PR–AUC, and ROC–AUC in some conditions, improvements were inconsistent and accompanied by greater variability. Graph neural network approaches did not provide consistent improvement and often produced lower decision-level performance despite increased model complexity. Given its strong performance, stability, interpretability, and computational efficiency, classical connectivity feature representation combined with anatomical encoding was selected as the primary modeling approach for all analyses in this study.

### 5 Supplementary Analysis: Effect of Trial Averaging Strategy

As described in the Methods, each electrode was recorded across approximately 400 trials spanning multiple language tasks, while clinical ground-truth labels were defined at the electrode level. Consequently, a key modeling consideration is how trial-level neural activity should be aggregated to produce a single prediction per electrode. We therefore compared two strategies for aggregating trial-level information into electrode-level predictions in order to determine whether preserving trial-to-trial variability improves classification performance.

**Post-hoc averaging (Trial-Level):** Connectivity features were extracted independently from each trial, and trial-level classifier predictions were averaged to produce electrode-level probabilities. This approach preserves trial-specific network variability and performs aggregation at the decision stage.

**Pre-hoc averaging (Averaged Signal):** Neural signals were first averaged across trials, and connectivity features were then extracted from the averaged signal. This approach reduces noise prior to feature extraction but removes trial-to-trial variability.

**Motor** For motor prediction, both strategies achieved strong performance, but trial-level modeling consistently yielded higher performance across metrics (Table S2, Table S3). Using classical connectivity features, trial-level modeling produced higher PR–AUC (0.755 vs 0.694), F1 score (0.753 vs 0.736), and balanced accuracy (0.872 vs 0.863), with similar ROC–AUC. Comparable reductions were observed for Node2Vec features. These findings indicate that while motor-critical electrodes are reliably identified under both approaches, preserving trial-level connectivity improves classification consistency.

Table S2: Effect of trial averaging on motor prediction using classical connectivity features.

| Model | AUC–PR | AUC–ROC | F1 | Balanced Acc |
|---|---|---|---|---|
| Trial-Level | 0.755 ± 0.191 | 0.929 ± 0.061 | 0.753 ± 0.143 | 0.872 ± 0.084 |
| Averaged | 0.694 ± 0.191 | 0.922 ± 0.059 | 0.736 ± 0.150 | 0.863 ± 0.083 |

Table S3: Effect of trial averaging on motor prediction using Node2Vec features.

| Model | AUC–PR | AUC–ROC | F1 | Balanced Acc |
|---|---|---|---|---|
| Trial-Level | 0.735 ± 0.179 | 0.929 ± 0.057 | 0.753 ± 0.143 | 0.872 ± 0.084 |
| Averaged | 0.710 ± 0.144 | 0.922 ± 0.074 | 0.710 ± 0.151 | 0.855 ± 0.082 |

**Speech Arrest** For speech arrest prediction, trial-level modeling consistently outperformed pre-hoc averaging across all performance metrics and connectivity representations (Table S4, Table S5). Using classical connectivity features, trial-level analysis achieved higher PR–AUC (0.550 vs 0.490), ROC–AUC (0.793 vs 0.770), F1 score (0.536 vs 0.442), and balanced accuracy (0.742 vs 0.665). Similar reductions were observed with Node2Vec features, including substantial decrease in F1 score (0.521 vs 0.365). These results indicate that trial-to-trial variability contains discriminative information that is lost when signals are averaged prior to feature extraction.

Table S4: Effect of trial averaging on speech arrest prediction using classical connectivity features.

| Model | AUC–PR | AUC–ROC | F1 | Balanced Acc |
|---|---|---|---|---|
| Trial-Level | 0.550 ± 0.196 | 0.793 ± 0.103 | 0.536 ± 0.233 | 0.742 ± 0.125 |
| Averaged | 0.490 ± 0.192 | 0.770 ± 0.093 | 0.442 ± 0.220 | 0.665 ± 0.128 |

Table S5: Effect of trial averaging on speech arrest prediction using Node2Vec features.

| Model | AUC–PR | AUC–ROC | F1 | Balanced Acc |
|---|---|---|---|---|
| Trial-Level | 0.569 ± 0.190 | 0.798 ± 0.101 | 0.521 ± 0.184 | 0.737 ± 0.111 |
| Averaged | 0.506 ± 0.202 | 0.784 ± 0.111 | 0.365 ± 0.192 | 0.625 ± 0.110 |

**Language** Language prediction showed smaller differences between strategies (Table S6, Table S7). Using classical connectivity features, trial-level modeling produced slightly higher PR–AUC, F1 score, and balanced accuracy. With Node2Vec features, averaged signals produced lower F1 score and balanced accuracy but slightly higher PR–AUC. Overall, differences were modest, indicating that trial averaging has less impact when discriminability is limited.

Table S6: Effect of trial averaging on language prediction using classical connectivity features.

| Model | AUC–PR | AUC–ROC | F1 | Balanced Acc |
|---|---|---|---|---|
| Trial-Level | 0.385 ± 0.167 | 0.761 ± 0.160 | 0.284 ± 0.189 | 0.675 ± 0.149 |
| Averaged | 0.365 ± 0.151 | 0.764 ± 0.159 | 0.274 ± 0.189 | 0.669 ± 0.168 |

Table S7: Effect of trial averaging on language prediction using Node2Vec features.

| Model | AUC–PR | AUC–ROC | F1 | Balanced Acc |
|---|---|---|---|---|
| Trial-Level | 0.432 ± 0.243 | 0.798 ± 0.158 | 0.314 ± 0.204 | 0.705 ± 0.154 |
| Averaged | 0.474 ± 0.291 | 0.791 ± 0.165 | 0.286 ± 0.223 | 0.649 ± 0.187 |

**Summary** Across functional domains and connectivity representations, trial-level modeling consistently achieved equal or higher performance than pre-hoc signal averaging, particularly for speech arrest and motor prediction. Improvements were observed not only in ranking metrics (AUC-PR and AUC-ROC) but also in classification metrics (F1 score and balanced accuracy), indicating more reliable identification of critical electrodes.

These findings show that preserving trial-specific connectivity patterns provides meaningful discriminative information that is diminished by early signal averaging. Based on this analysis, all primary models and results presented in this study were trained using trial-level connectivity features with post-hoc averaging of predictions.

#### 6 Supplementary Analysis: Effect of Classifier Choice

To determine how classifier selection influences prediction performance, we compared multiple linear, nonlinear, and ensemble classifiers using the identical leave-one-subject-out framework and feature representation employed throughout the primary analyses (anatomical region encoding combined with connectivity features derived from the production-aligned window). This comparison isolates the effect of classifier formulation while holding all other aspects of the analysis constant. We evaluated the following classifiers:

**Support Vector Machine (Linear and RBF kernels)** Support vector machine classifiers were trained using both linear and radial basis function (RBF) kernels. Balanced class weights were applied to compensate for class imbalance by weighting classes inversely to their frequency.

**Logistic Regression** A probabilistic linear classifier trained using the SAGA solver with L2 regularization ($C = 0.5$) and balanced class weights.

**Random Forest** An ensemble of 100 decision trees trained using bootstrap aggregation. Tree depth was limited to 15, with minimum samples per split and leaf set to 5 and 2 respectively.

**Decision Tree** A single decision tree with maximum depth of 10 and balanced class weights.

**Multilayer Perceptron (MLP)** A two-layer neural network (64 and 32 hidden units) with ReLU activations, trained using Adam optimization and early stopping.

##### Motor

Motor prediction results are shown in Figure S11 and were uniformly high across classifiers. Linear SVM and logistic regression achieved the strongest overall performance. The MLP and decision tree produced similar results to the linear models, whereas random forest showed reduced F1 score and balanced accuracy despite comparable ROC–AUC. Overall, performance differences between classifiers were small, consistent with the strong anatomical specificity of motor cortex reflected in the feature representation.

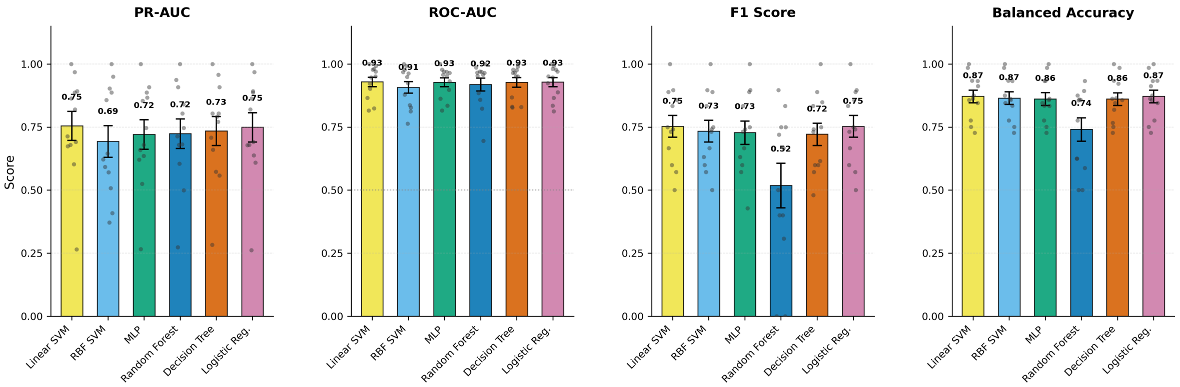


Figure S11: Classifier comparison for Motor prediction. Bars show mean ± SEM across LOSO folds; gray dots indicate individual fold values.

#### Speech Arrest

Classifier performance for speech arrest prediction is summarized in Figure S12. Linear SVM and logistic regression yielded close performance across all metrics, reflecting the suitability of linear decision boundaries for this feature representation. The RBF SVM achieved comparable ROC–AUC but showed lower F1 score and balanced accuracy, indicating less reliable thresholded classification despite similar ranking ability. Tree-based models and the MLP achieved similar or slightly higher ROC–AUC values but did not consistently improve F1 score or balanced accuracy relative to the linear models. These results suggest that increased model complexity did not translate into more stable electrode-level classification for speech arrest.

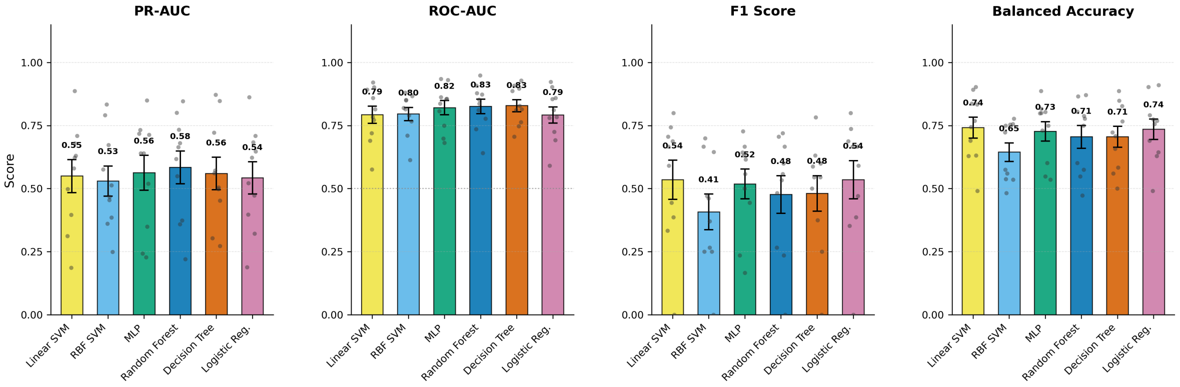


Figure S12: Classifier comparison for Speech Arrest prediction.

#### Language

Language prediction performance is summarized in Figure S13. Overall performance was modest across all classifiers. Linear SVM and logistic regression achieved the strongest and most consistent results (PR–AUC $0.385 \pm 0.167$ and $0.384 \pm 0.167$, ROC–AUC $0.761 \pm 0.160$ and $0.768 \pm 0.150$, respectively), with similar F1 scores and balanced accuracy. Other classifiers did not improve performance. The RBF SVM showed lower discrimination across both PR–AUC and ROC–AUC, while tree-based models and the MLP achieved slightly lower AUC values and lower F1 scores and balanced accuracy.

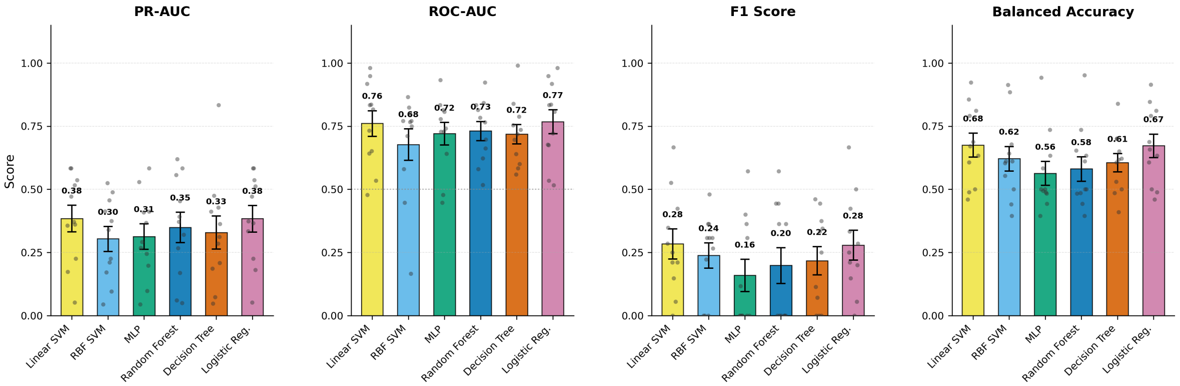


Figure S13: Classifier comparison for Language prediction.

#### Summary

Linear SVM consistently achieved strong and stable performance across all metrics, matching or exceeding more complex classifiers in F1 score and balanced accuracy while maintaining competitive ROC–AUC and PR–AUC. In contrast, nonlinear classifiers did not provide consistent improvements and in some cases showed reduced classification stability. Based on these results and given its strong overall performance, stability, robustness to high-dimensional feature spaces, and widespread use in neurophysiological decoding studies, the linear SVM was selected as the primary classifier for all main analyses presented in this paper.

### 7 Supplementary Analysis: Effect of Functional Definition

Additional threshold-dependent metrics, including positive predictive value (PPV), negative predictive value (NPV), sensitivity, and specificity, are summarized in Table S8.

Table S8: Threshold-dependent performance metrics across models (mean values across subjects).

| Model | PPV | NPV | Sens | Spec | F1 | Bal Acc |
|---|---|---|---|---|---|---|
| Motor | 0.741 | 0.964 | 0.817 | 0.926 | 0.753 | 0.872 |
| Speech Arrest | 0.471 | 0.912 | 0.682 | 0.802 | 0.536 | 0.742 |
| Language | 0.215 | 0.928 | 0.672 | 0.679 | 0.284 | 0.675 |
| Combined (Single Model) | 0.522 | 0.849 | 0.840 | 0.567 | 0.612 | 0.704 |
| Combined (Max-Fusion) | 0.660 | 0.882 | 0.844 | 0.620 | 0.686 | 0.732 |

To provide a detailed view of classifier performance beyond summary metrics, we examined receiver operating characteristic (ROC) and precision–recall (PR) curves for each ground-truth label definition. Supplementary Figures S14–S18 show ROC and PR curves across leave-one-subject-out folds for motor, speech arrest, language, Combined (Single Model and Max-Fusion) definitions. Thin gray lines indicate individual folds, and the thick colored line indicates the mean performance across folds.

Consistent with the summary metrics reported in Table 3, motor-critical cortex exhibited the strongest discrimination, with ROC and PR curves demonstrating consistently high sensitivity and precision across thresholds. Speech arrest showed intermediate performance, with reliable separation between critical and non-critical electrodes. Language-critical cortex exhibited lower precision and greater variability across folds, reflecting increased biological heterogeneity.

Combined Single Model and Max-Fusion strategies maintained strong ROC performance but showed lower PR-AUC lift due to increased prevalence of positive electrodes under these definitions. These curves illustrate the trade-off between sensitivity and precision across different labeling strategies and confirm the robustness of the reported performance metrics.

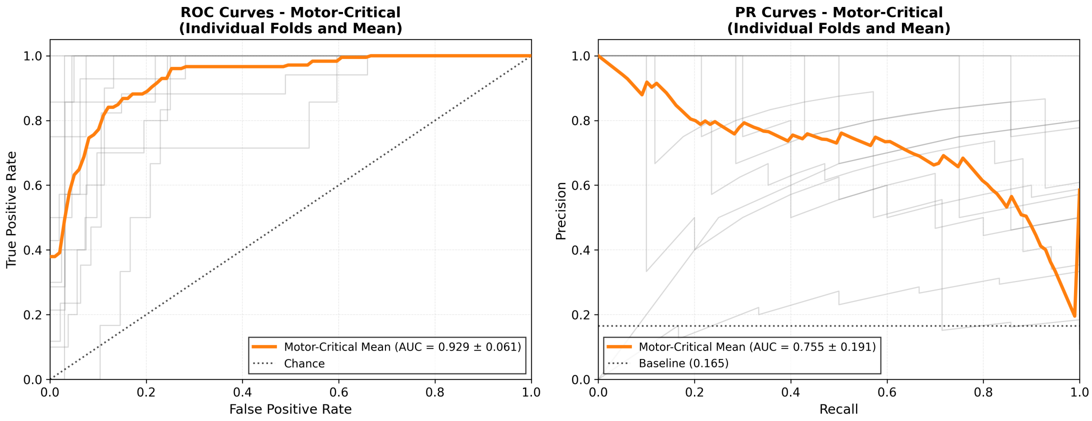


Figure S14: ROC and Precision–Recall curves for motor prediction. Thin gray lines indicate individual leave-one-subject-out folds, and the thick line indicates the mean performance. The dotted line indicates chance performance. Motor-critical cortex showed consistently high sensitivity and precision across folds.

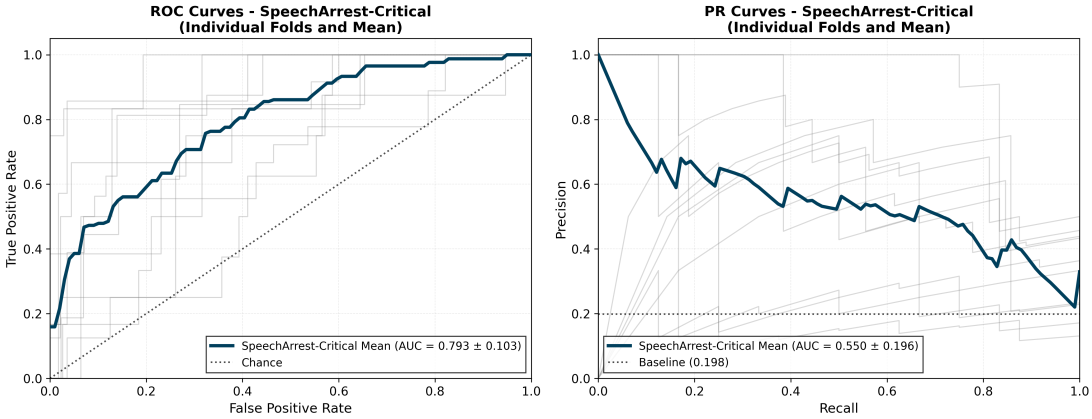


Figure S15: ROC and Precision–Recall curves for speech arrest prediction.

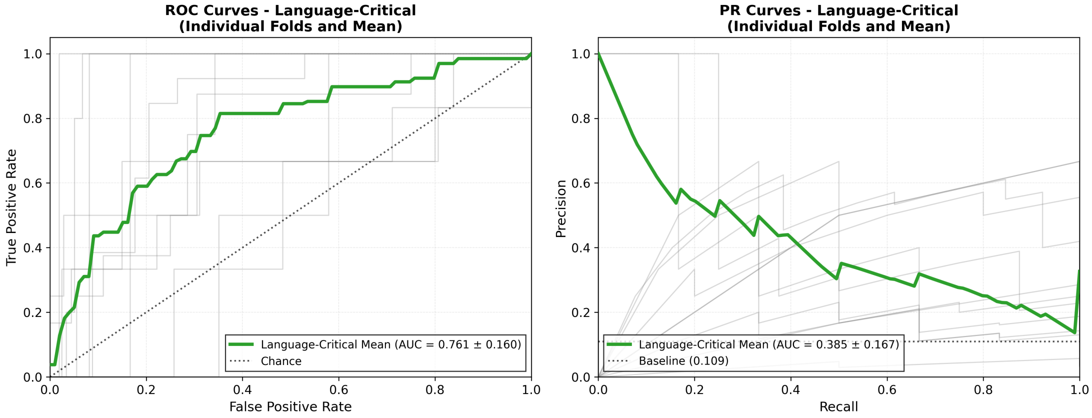


Figure S16: ROC and Precision–Recall curves for language prediction. Greater variability and lower precision reflect the distributed organization of language networks.

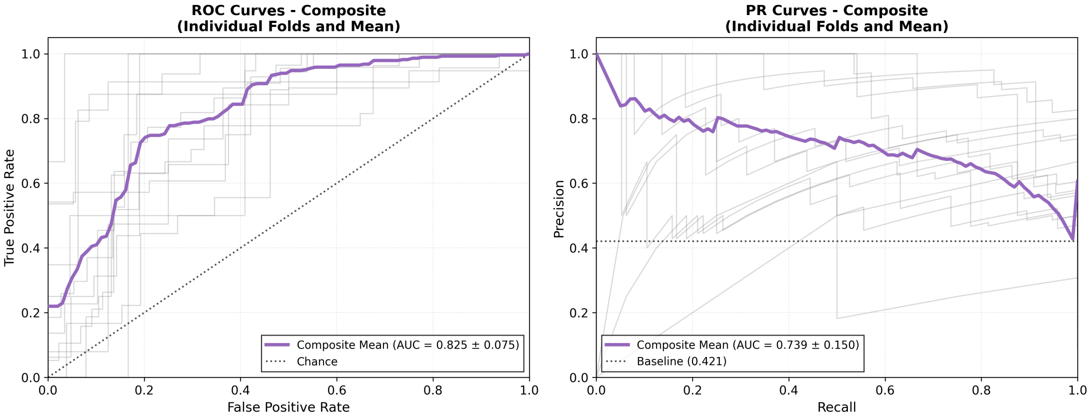


Figure S17: ROC and Precision–Recall curves for the Combined (Single Model) strategy.

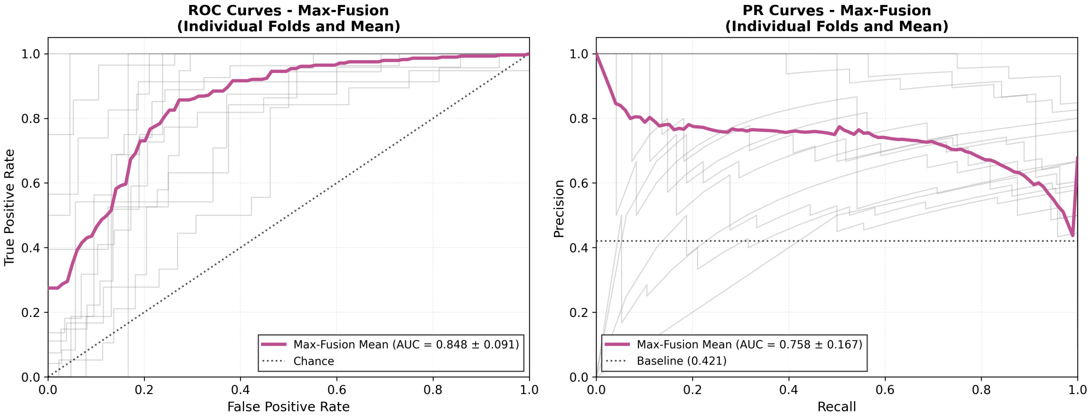


Figure S18: ROC and Precision–Recall curves for the Combined (Max-Fusion) strategy.

### 8 Supplementary Analysis: Full Task Composition Performance

To provide complete quantitative detail for the task-composition analysis (Figure 2), we report full performance tables for all task configurations, including single tasks, task pairs, task trios, and the full task set, separately for motor-, speech arrest-, and language-critical definitions.

Table S1: Full task-composition performance for motor-critical prediction.

| Task Combination | ROC–AUC | PR–AUC | Prev | PR Lift | Avg Score |
|---|---|---|---|---|---|
| **Single Tasks** | | | | | |
| PicN | 0.935 ± 0.055 | 0.775 ± 0.175 | 0.153 | 5.061 | 2.998 |
| AudRep | 0.931 ± 0.061 | 0.749 ± 0.194 | 0.153 | 4.894 | 2.913 |
| VisRead | 0.928 ± 0.063 | 0.741 ± 0.197 | 0.153 | 4.840 | 2.884 |
| AudN | 0.921 ± 0.067 | 0.700 ± 0.204 | 0.153 | 4.589 | 2.755 |
| SenComp | 0.937 ± 0.061 | 0.785 ± 0.181 | 0.183 | 4.297 | 2.617 |
| **Task Pairs** | | | | | |
| AudRep+PicN | 0.927 ± 0.067 | 0.754 ± 0.189 | 0.153 | 4.928 | 2.928 |
| PicN+VisRead | 0.933 ± 0.057 | 0.750 ± 0.191 | 0.153 | 4.901 | 2.917 |
| AudRep+VisRead | 0.932 ± 0.059 | 0.747 ± 0.197 | 0.153 | 4.876 | 2.904 |
| PicN+SenComp | 0.933 ± 0.058 | 0.781 ± 0.167 | 0.165 | 4.722 | 2.827 |
| AudN+PicN | 0.928 ± 0.062 | 0.762 ± 0.176 | 0.165 | 4.603 | 2.766 |
| AudRep+SenComp | 0.927 ± 0.066 | 0.758 ± 0.196 | 0.165 | 4.579 | 2.753 |
| SenComp+VisRead | 0.930 ± 0.058 | 0.753 ± 0.191 | 0.165 | 4.551 | 2.740 |
| AudN+AudRep | 0.928 ± 0.061 | 0.747 ± 0.193 | 0.165 | 4.514 | 2.721 |
| AudN+VisRead | 0.925 ± 0.064 | 0.737 ± 0.196 | 0.165 | 4.457 | 2.691 |
| AudN+SenComp | 0.927 ± 0.062 | 0.730 ± 0.216 | 0.165 | 4.413 | 2.670 |
| **Task Trios** | | | | | |
| AudRep+PicN+VisRead | 0.931 ± 0.059 | 0.753 ± 0.192 | 0.153 | 4.916 | 2.923 |
| AudRep+PicN+SenComp | 0.930 ± 0.064 | 0.767 ± 0.186 | 0.165 | 4.633 | 2.782 |
| AudN+AudRep+PicN | 0.930 ± 0.063 | 0.763 ± 0.189 | 0.165 | 4.612 | 2.771 |
| PicN+SenComp+VisRead | 0.935 ± 0.052 | 0.756 ± 0.185 | 0.165 | 4.568 | 2.752 |
| AudRep+SenComp+VisRead | 0.931 ± 0.058 | 0.756 ± 0.192 | 0.165 | 4.569 | 2.750 |
| AudN+PicN+SenComp | 0.931 ± 0.058 | 0.756 ± 0.185 | 0.165 | 4.568 | 2.749 |
| AudN+AudRep+SenComp | 0.928 ± 0.064 | 0.756 ± 0.197 | 0.165 | 4.570 | 2.749 |
| AudN+AudRep+VisRead | 0.930 ± 0.059 | 0.749 ± 0.193 | 0.165 | 4.524 | 2.727 |
| AudN+PicN+VisRead | 0.927 ± 0.063 | 0.746 ± 0.188 | 0.165 | 4.507 | 2.717 |
| AudN+SenComp+VisRead | 0.926 ± 0.065 | 0.742 ± 0.201 | 0.165 | 4.482 | 2.704 |
| **All Tasks** | | | | | |
| All Tasks | 0.929 ± 0.061 | 0.755 ± 0.191 | 0.165 | 4.561 | 2.745 |

Table S2: Full task-composition performance for speech arrest-critical prediction.

| Task Combination | ROC–AUC | PR–AUC | Prev | PR Lift | Avg Score |
|---|---|---|---|---|---|
| **Single Tasks** | | | | | |
| AudRep | 0.786 ± 0.096 | 0.536 ± 0.198 | 0.179 | 2.992 | 1.889 |
| AudN | 0.833 ± 0.052 | 0.585 ± 0.171 | 0.209 | 2.797 | 1.815 |
| PicN | 0.796 ± 0.103 | 0.499 ± 0.211 | 0.179 | 2.785 | 1.791 |
| VisRead | 0.797 ± 0.093 | 0.495 ± 0.199 | 0.179 | 2.767 | 1.782 |
| SenComp | 0.672 ± 0.123 | 0.360 ± 0.149 | 0.198 | 1.816 | 1.244 |
| **Task Pairs** | | | | | |
| AudRep+PicN | 0.789 ± 0.101 | 0.512 ± 0.207 | 0.179 | 2.860 | 1.824 |
| AudRep+VisRead | 0.790 ± 0.090 | 0.502 ± 0.196 | 0.179 | 2.806 | 1.798 |
| PicN+VisRead | 0.797 ± 0.097 | 0.501 ± 0.205 | 0.179 | 2.797 | 1.797 |
| AudN+AudRep | 0.794 ± 0.092 | 0.555 ± 0.194 | 0.198 | 2.798 | 1.796 |
| AudN+PicN | 0.799 ± 0.099 | 0.541 ± 0.203 | 0.198 | 2.729 | 1.764 |
| AudN+SenComp | 0.792 ± 0.111 | 0.542 ± 0.196 | 0.198 | 2.731 | 1.761 |
| AudRep+SenComp | 0.782 ± 0.110 | 0.539 ± 0.192 | 0.198 | 2.718 | 1.750 |
| AudN+VisRead | 0.798 ± 0.089 | 0.532 ± 0.193 | 0.198 | 2.681 | 1.740 |
| PicN+SenComp | 0.787 ± 0.111 | 0.531 ± 0.198 | 0.198 | 2.676 | 1.732 |
| SenComp+VisRead | 0.786 ± 0.107 | 0.522 ± 0.192 | 0.198 | 2.629 | 1.707 |
| **Task Trios** | | | | | |
| AudRep+PicN+VisRead | 0.795 ± 0.096 | 0.514 ± 0.197 | 0.179 | 2.868 | 1.832 |
| AudN+AudRep+SenComp | 0.795 ± 0.100 | 0.560 ± 0.201 | 0.198 | 2.822 | 1.809 |
| AudN+PicN+SenComp | 0.796 ± 0.104 | 0.548 ± 0.199 | 0.198 | 2.763 | 1.779 |
| AudN+AudRep+PicN | 0.793 ± 0.102 | 0.546 ± 0.204 | 0.198 | 2.753 | 1.773 |
| AudN+SenComp+VisRead | 0.796 ± 0.096 | 0.541 ± 0.190 | 0.198 | 2.727 | 1.761 |
| AudRep+PicN+SenComp | 0.787 ± 0.107 | 0.541 ± 0.198 | 0.198 | 2.728 | 1.758 |
| AudRep+SenComp+VisRead | 0.788 ± 0.100 | 0.539 ± 0.191 | 0.198 | 2.715 | 1.752 |
| AudN+AudRep+VisRead | 0.791 ± 0.094 | 0.536 ± 0.196 | 0.198 | 2.699 | 1.745 |
| AudN+PicN+VisRead | 0.794 ± 0.099 | 0.527 ± 0.202 | 0.198 | 2.654 | 1.724 |
| PicN+SenComp+VisRead | 0.791 ± 0.104 | 0.523 ± 0.202 | 0.198 | 2.636 | 1.714 |
| **All Tasks** | | | | | |
| All Tasks | 0.793 ± 0.103 | 0.550 ± 0.196 | 0.198 | 2.772 | 1.783 |

Table S3: Full task-composition performance for language-critical prediction.

| Task Combination | ROC–AUC | PR–AUC | Prev | PR Lift | Avg Score |
|---|---|---|---|---|---|
| **Single Tasks** | | | | | |
| SenComp | 0.752 ± 0.175 | 0.511 ± 0.165 | 0.108 | 4.741 | 2.746 |
| AudRep | 0.765 ± 0.167 | 0.385 ± 0.173 | 0.107 | 3.586 | 2.175 |
| VisRead | 0.772 ± 0.158 | 0.380 ± 0.166 | 0.107 | 3.543 | 2.157 |
| AudN | 0.751 ± 0.142 | 0.404 ± 0.196 | 0.114 | 3.553 | 2.152 |
| PicN | 0.757 ± 0.159 | 0.342 ± 0.151 | 0.105 | 3.259 | 2.008 |
| **Task Pairs** | | | | | |
| AudRep+PicN | 0.763 ± 0.167 | 0.402 ± 0.213 | 0.107 | 3.748 | 2.256 |
| AudN+SenComp | 0.781 ± 0.144 | 0.405 ± 0.196 | 0.109 | 3.704 | 2.242 |
| AudRep+VisRead | 0.772 ± 0.163 | 0.396 ± 0.177 | 0.107 | 3.689 | 2.230 |
| AudRep+SenComp | 0.763 ± 0.162 | 0.394 ± 0.170 | 0.109 | 3.603 | 2.183 |
| AudN+VisRead | 0.769 ± 0.150 | 0.391 ± 0.163 | 0.109 | 3.576 | 2.172 |
| AudN+AudRep | 0.753 ± 0.165 | 0.385 ± 0.165 | 0.109 | 3.521 | 2.137 |
| AudN+PicN | 0.760 ± 0.149 | 0.383 ± 0.161 | 0.109 | 3.499 | 2.129 |
| SenComp+VisRead | 0.774 ± 0.149 | 0.376 ± 0.163 | 0.109 | 3.440 | 2.107 |
| PicN+VisRead | 0.766 ± 0.158 | 0.370 ± 0.166 | 0.107 | 3.446 | 2.106 |
| PicN+SenComp | 0.763 ± 0.154 | 0.376 ± 0.174 | 0.109 | 3.442 | 2.102 |
| **Task Trios** | | | | | |
| AudRep+PicN+SenComp | 0.762 ± 0.163 | 0.428 ± 0.240 | 0.109 | 3.913 | 2.338 |
| AudN+AudRep+SenComp | 0.766 ± 0.160 | 0.419 ± 0.213 | 0.109 | 3.830 | 2.298 |
| AudRep+SenComp+VisRead | 0.765 ± 0.162 | 0.411 ± 0.209 | 0.109 | 3.761 | 2.263 |
| AudN+AudRep+VisRead | 0.763 ± 0.156 | 0.394 ± 0.168 | 0.109 | 3.600 | 2.181 |
| AudN+SenComp+VisRead | 0.776 ± 0.151 | 0.389 ± 0.168 | 0.109 | 3.553 | 2.164 |
| AudN+PicN+SenComp | 0.767 ± 0.152 | 0.388 ± 0.168 | 0.109 | 3.546 | 2.156 |
| AudRep+PicN+VisRead | 0.766 ± 0.163 | 0.381 ± 0.172 | 0.107 | 3.546 | 2.156 |
| AudN+AudRep+PicN | 0.756 ± 0.159 | 0.385 ± 0.165 | 0.109 | 3.523 | 2.140 |
| PicN+SenComp+VisRead | 0.765 ± 0.153 | 0.383 ± 0.165 | 0.109 | 3.505 | 2.135 |
| AudN+PicN+VisRead | 0.756 ± 0.156 | 0.380 ± 0.161 | 0.109 | 3.472 | 2.114 |
| **All Tasks** | | | | | |
| All Tasks | 0.761 ± 0.160 | 0.385 ± 0.167 | 0.109 | 3.519 | 2.140 |

#### 9 Supplementary Analysis: Effect of Number of Trials

To provide detailed quantitative support for the trial-count analysis presented in the main text (Figure 3), we report full performance metrics across all evaluated trial counts for auditory repetition and picture naming tasks. Tables S4–S6 summarize mean performance and standard deviation across random seeds for motor, speech arrest, and language criticality definitions, respectively.

Table S4: Performance across trial counts for motor prediction.

| Trials | ROC–AUC | PR–AUC | F1 | Balanced Acc |
|---|---|---|---|---|
| *Picture Naming* | | | | |
| 1 | 0.930 ± 0.007 | 0.735 ± 0.015 | 0.726 ± 0.014 | 0.858 ± 0.010 |
| 5 | 0.928 ± 0.004 | 0.758 ± 0.009 | 0.717 ± 0.025 | 0.856 ± 0.011 |
| 10 | 0.935 ± 0.007 | 0.770 ± 0.013 | 0.733 ± 0.005 | 0.863 ± 0.003 |
| 25 | 0.932 ± 0.007 | 0.769 ± 0.011 | 0.735 ± 0.002 | 0.864 ± 0.001 |
| 50 | 0.933 ± 0.008 | 0.765 ± 0.010 | 0.739 ± 0.002 | 0.866 ± 0.001 |
| 75 | 0.932 ± 0.002 | 0.771 ± 0.008 | 0.739 ± 0.003 | 0.866 ± 0.001 |
| 100 | 0.935 | 0.775 | 0.740 | 0.866 |
| *Auditory Repetition* | | | | |
| 1 | 0.927 ± 0.011 | 0.735 ± 0.038 | 0.736 ± 0.004 | 0.862 ± 0.003 |
| 5 | 0.929 ± 0.007 | 0.761 ± 0.015 | 0.738 ± 0.003 | 0.865 ± 0.002 |
| 10 | 0.928 ± 0.010 | 0.752 ± 0.019 | 0.732 ± 0.012 | 0.861 ± 0.007 |
| 25 | 0.932 ± 0.008 | 0.756 ± 0.015 | 0.740 ± 0.000 | 0.866 ± 0.000 |
| 50 | 0.932 ± 0.008 | 0.753 ± 0.007 | 0.740 ± 0.000 | 0.866 ± 0.000 |
| 75 | 0.931 ± 0.006 | 0.751 ± 0.008 | 0.740 ± 0.000 | 0.866 ± 0.000 |
| 100 | 0.931 | 0.749 | 0.740 | 0.866 |

Table S5: Performance across trial counts for speech arrest prediction.

| Trials | ROC–AUC | PR–AUC | F1 | Balanced Acc |
|---|---|---|---|---|
| | | *Picture Naming* | | |
| 1 | 0.773 ± 0.009 | 0.461 ± 0.017 | 0.474 ± 0.014 | 0.730 ± 0.012 |
| 5 | 0.791 ± 0.010 | 0.515 ± 0.026 | 0.478 ± 0.016 | 0.713 ± 0.025 |
| 10 | 0.793 ± 0.007 | 0.504 ± 0.025 | 0.474 ± 0.025 | 0.712 ± 0.028 |
| 25 | 0.793 ± 0.004 | 0.502 ± 0.006 | 0.473 ± 0.037 | 0.709 ± 0.018 |
| 50 | 0.801 ± 0.003 | 0.520 ± 0.012 | 0.475 ± 0.020 | 0.706 ± 0.007 |
| 75 | 0.798 ± 0.005 | 0.503 ± 0.007 | 0.432 ± 0.020 | 0.687 ± 0.016 |
| 100 | 0.796 | 0.499 | 0.443 | 0.697 |
| | | *Auditory Repetition* | | |
| 1 | 0.784 ± 0.010 | 0.497 ± 0.027 | 0.469 ± 0.011 | 0.724 ± 0.014 |
| 5 | 0.791 ± 0.006 | 0.519 ± 0.019 | 0.483 ± 0.021 | 0.710 ± 0.024 |
| 10 | 0.788 ± 0.004 | 0.523 ± 0.020 | 0.483 ± 0.012 | 0.713 ± 0.009 |
| 25 | 0.787 ± 0.007 | 0.525 ± 0.012 | 0.463 ± 0.012 | 0.702 ± 0.008 |
| 50 | 0.788 ± 0.005 | 0.536 ± 0.015 | 0.475 ± 0.007 | 0.710 ± 0.006 |
| 75 | 0.787 ± 0.001 | 0.539 ± 0.007 | 0.475 ± 0.009 | 0.710 ± 0.008 |
| 100 | 0.786 | 0.536 | 0.491 | 0.722 |

Table S6: Performance across trial counts for language prediction.

| Trials | ROC–AUC | PR–AUC | F1 | Balanced Acc |
|---|---|---|---|---|
| | | *Picture Naming* | | |
| 1 | 0.747 ± 0.006 | 0.365 ± 0.019 | 0.239 ± 0.024 | 0.631 ± 0.017 |
| 5 | 0.752 ± 0.005 | 0.352 ± 0.031 | 0.210 ± 0.031 | 0.621 ± 0.016 |
| 10 | 0.759 ± 0.006 | 0.367 ± 0.018 | 0.228 ± 0.014 | 0.631 ± 0.005 |
| 25 | 0.758 ± 0.005 | 0.369 ± 0.028 | 0.234 ± 0.022 | 0.647 ± 0.019 |
| 50 | 0.756 ± 0.002 | 0.355 ± 0.023 | 0.226 ± 0.021 | 0.637 ± 0.020 |
| 75 | 0.756 ± 0.003 | 0.350 ± 0.017 | 0.225 ± 0.015 | 0.634 ± 0.014 |
| 100 | 0.757 | 0.342 | 0.242 | 0.658 |
| | | *Auditory Repetition* | | |
| 1 | 0.753 ± 0.021 | 0.372 ± 0.053 | 0.256 ± 0.015 | 0.673 ± 0.023 |
| 5 | 0.772 ± 0.004 | 0.400 ± 0.014 | 0.261 ± 0.009 | 0.675 ± 0.013 |
| 10 | 0.774 ± 0.004 | 0.404 ± 0.020 | 0.261 ± 0.009 | 0.673 ± 0.004 |
| 25 | 0.770 ± 0.007 | 0.396 ± 0.030 | 0.271 ± 0.006 | 0.679 ± 0.004 |
| 50 | 0.768 ± 0.002 | 0.400 ± 0.024 | 0.274 ± 0.008 | 0.681 ± 0.006 |
| 75 | 0.768 ± 0.003 | 0.396 ± 0.016 | 0.276 ± 0.004 | 0.682 ± 0.004 |
| 100 | 0.765 | 0.385 | 0.274 | 0.683 |
